# High-quality haploid genomes corroborate 29 chromosomes and highly conserved synteny of genes in *Hyles* hawkmoths (Lepidoptera: Sphingidae)

**DOI:** 10.1101/2022.04.08.487644

**Authors:** Anna K. Hundsdoerfer, Tilman Schell, Franziska Patzold, Charlotte J. Wright, Atsuo Yoshido, František Marec, Hana Daneck, Sylke Winkler, Carola Greve, Lars Podsiadlowski, Michael Hiller, Martin Pippel

**Affiliations:** Senckenberg Natural History Collections Dresden, Königsbrücker Landstr. 159, 01109 Dresden, Germany; LOEWE-Centre for Translational Biodiversity Genomics (LOEWE-TBG), Frankfurt am Main, Germany; Tree of Life, Wellcome Sanger Institute, Cambridge, CB10 1SA, UK; Biology Centre of the Czech Academy of Sciences, Institute of Entomology, Branišovská 31, 370 05 České Budějovice, Czech Republic; Max Planck Institute of Molecular Cell Biology and Genetics, Pfotenhauerstraße 108, 01307 Dresden, Germany; Centre for Molecular Biodiversity Research, Leibniz Institute for the Analysis of Biodiversity Change, Adenauerallee 127, 53113 Bonn, Germany; Center for Systems Biology Dresden, Pfotenhauerstr. 108, 01307 Dresden, Germany

**Keywords:** karyotype, chromosome-level scaffolding, wing pattern genes, *optix*, *Wnt*, *cortex*, *aristaless*, *distal-less*, *P* supergene

## Abstract

**Background:** Morphological and traditional genetic studies of the young Pliocene genus *Hyles* have led to the understanding that despite its importance for taxonomy, phenotypic similarity of wing patterns does not correlate with phylogenetic relationship. To gain insights into various aspects of speciation in the Spurge Hawkmoth (*Hyles euphorbiae*), we assembled a chromosome-level genome and investigated some of its characteristics.

**Results:** The genome of a male *H. euphorbiae* was sequenced using PacBio and Hi-C data, yielding a 504 Mb assembly (scaffold N50 of 18.2 Mb) with 99.9% of data represented by the 29 largest scaffolds forming the haploid chromosome set. Consistent with this, FISH analysis of the karyotype revealed *n* = 29 chromosomes and a WZ/ZZ (female/male) sex chromosome system. Estimates of chromosome length based on the karyotype image provided an additional quality metric of assembled chromosome size. Rescaffolding the published male *H. vespertilio* genome resulted in a high-quality assembly (651 Mb, scaffold N50 of 22 Mb) with 98% of sequence data in the 29 chromosomes. The larger genome size of *H. vespertilio* (average 1C DNA value of 562 Mb) was accompanied by a proportional increase in repeats from 45% in *H. euphorbiae* (measured as 472 Mb) to almost 55% in *H. vespertilio*. Several wing pattern genes were found on the same chromosomes in the two species, with varying amounts and positions of repetitive elements and inversions possibly corrupting their function.

**Conclusions:** Our two-fold comparative genomics approach revealed high gene synteny of the *Hyles* genomes to other Sphingidae and high correspondence to intact Merian elements, the ancestral linkage groups of Lepidoptera, with the exception of three simple fusion events. We propose a standardized approach for genome taxonomy using nucleotide homology via scaffold chaining as the primary tool combined with Oxford plots based on Merian elements to infer and visualize directionality of chromosomal rearrangements. The identification of wing pattern genes promises future understanding of the evolution of forewing patterns in the genus *Hyles*, although further sequencing data from more individuals are needed. The genomic data obtained provide additional reliable references for further comparative studies in hawkmoths (Sphingidae).

## Background

The genus *Hyles* comprises about 30 species, is distributed worldwide and, with an estimated divergence time of 5.7-9.3 million years (Mya) [1], is rather young from an evolutionary perspective [2]. Some species are still in the process of speciation, and others hybridize because reproductive barriers are still low. These processes are difficult to discern in *Hyles euphorbiae*, which was previously comprised of five separate species [3]. The Spurge Hawkmoth, *Hyles euphorbiae* is a charismatic Palearctic species with large, colorful, and polymorphic larvae and camouflaged, heavy adults with strong flight abilities (Fig. 1a). Despite their aposematic coloration, the larvae do not sequester the toxic spurge diterpene esters [4] from their food plants. Instead, they protect themselves by spewing the plant slurry that has not yet been detoxified. Larval specialization on toxic host plants of the genus *Euphorbia* has evolved twice independently within the genus *Hyles* [5, 6]. The impressively high morphological variability of larvae has complicated the taxonomy of *H. euphorbiae* by contributing to an overestimation of species diversity (overview in Hundsdoerfer et al. [3]). Similarly, the high intraspecific diversity of mitochondrial marker genes bedeviled the reconstruction of the molecular phylogeny of the formerly five species (synonymized with *H. euphorbiae* in 2019 [5]), but provided valuable resolution for phylogeography [7]. Although some larval patterns are correlated with geography [7] and are thus expected to be based on underlying genetic diversity causing phenotypic variability, others appear to be environmentally driven by phenotypic plasticity.

**Fig. 1.**
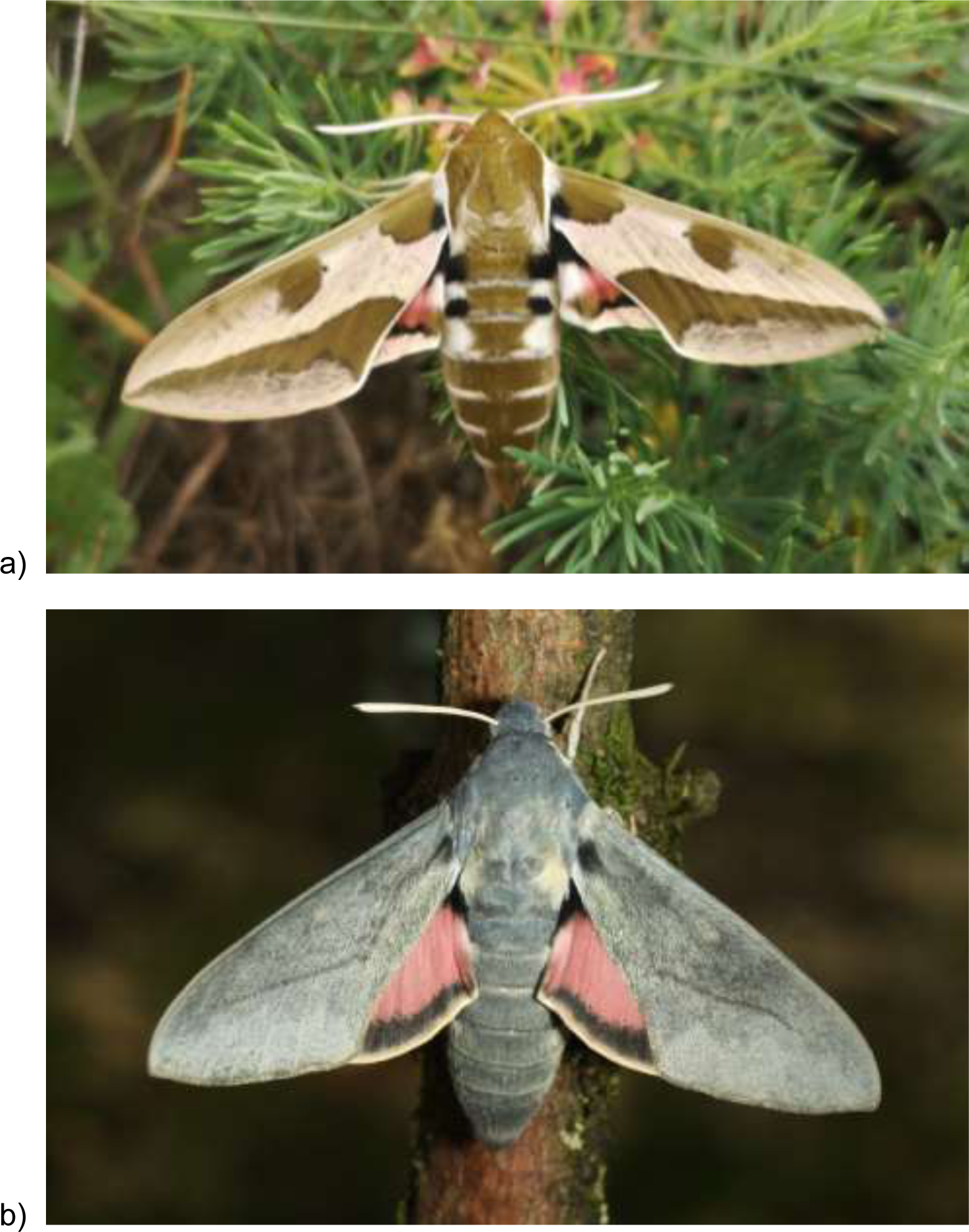
**a)** A male of the Spurge Hawkmoth (*Hyles euphorbiae*) collected near Berbisdorf, Meissen, Germany (photo: AKH); **b)** A female of the Bat Hawkmoth (*Hyles vespertilio*) photographed by Jean Haxaire in l’Alpes d’Huez, France (copyright and courtesy of Jean Haxaire [8]).

Early studies have shown that similarity in wing patterns is not based on phylogenetic affiliation in this genus [1, 2]. Seven basic wing patterns are observed in the Central Palearctic *Hyles* species, which neither correlate with currently defined species, nor reflect phylogenetic relationships within the genus. One very prominent wing pattern shows many white stripes on brown forewings, a set of shared character states [9] that originally led to the assumption that the Palearctic *H. livornica* and the Australian *H. livornicoides* were conspecific with the New World species *H. lineata* (e.g., Haruta, 1994). After their recognition as three species, they were still considered to be a distinct subgenus ‘Danneria’ [10], but this has been refuted as a clade by previous molecular phylogenetic work [2]. The species group in very different parts of the tree and do not form a tight cluster of cryptic species. Another prominent wing pattern is one of dark brown spots and stripes on a lighter, cream-colored background, corresponding to the typical forewing pattern of *H. euphorbiae*, and thus the group included numerous species with a similar wing pattern (or slight variations thereof) [9]. However, again, some of these, e.g., *H. centralasiae*, *H. nicaea,* and *H. euphorbiae* were not found to have a close molecular phylogenetic relationship [11]. The species *H. vespertilio* (Fig. 1b), the genome of which was recently published [12], has a forewing that lacks the pattern elements described. The forewings of this species almost completely lack a wing pattern [12], raising the possibility that the gene(s) controlling wing pattern formation are inactive, making genomic comparisons with this species particularly intriguing.

Interspecific differences in forewing patterns within the genus *Hyles* should be based on detectable genetic differences, as patterns are stable within species in the well-separated, oldest Neotropical (and Nearctic) taxa [2]. In the Palearctic, incomplete lineage sorting and ongoing hybridization impede such insights, justifying ongoing systematic, phylogenetic and taxonomic research (e.g., Patzold et al. [13]).

Genomic data promise insights into numerous aspects of speciation, including ongoing gene flow, food plant utilization, and the genetic basis for morphological differentiation. Therefore, we sequenced and examined the genome of *Hyles euphorbiae*. Numerous Sphingidae genomes are currently being published (e.g. *Hyles vespertilio*, Macroglossinae, by Pippel et al. [12], *Mimas tiliae*, Smerinthinae, by Boyes et al. [14], *Laothoe populi*, Smerinthinae [15]). This wealth of data will provide insights into the evolution of wing patterns in hawkmoths. Pioneering studies of the genes underlying the wing pattern of Lepidoptera have been performed on the family Nymphalidae, facilitated by a classical model describing wing-pattern elements (e.g. eyespots, bands) in an idealized ‘nymphalid groundplan’ [16, 17]. The genetic induction of colored scales of pattern elements on Lepidoptera wings has been well-studied in several genera of the family Nymphalidae, namely *Bicyclus* [18, 19], *Heliconius* [20]*, Junonia* [21], and *Vanessa* [22]. It has been previously proposed that the evolution of lepidopteran wing pattern stripes occurred through the repeated gain, loss, and modification of only a handful of serially repeated elements, arranged in a modular architecture with narrow stretches of the genome associated with specific differences in color and pattern [23]. In addition, phenotypic plasticity in variation of colors and certain color patterns of species is expected to be primarily driven by biotic and abiotic factors [24] acting as a selection pressure on individuals with differential expression of genotypes [18, 25]. Strong selection pressure can limit polymorphism, keeping the best-protected wing pattern, but is has been shown that alternative adaptive phenotypes can be maintained by heterozygote genotypes of the wing-pattern *P* supergene in natural populations of *Heliconius numata* [26]. The *P* supergene represents a non-recombining part of chromosome 15 consisting of three stretches with co-adaptive loci (P_1_-P_3_ of sizes 400 kb, 200 kb and 1150 kb) that are kept in close linkage by polymorphic inversions [27].

*Optix* is a single-exon gene in *Heliconius* [20, 28] encoding a homeobox transcription factor that regulates the production of ommochrome pigments [29] and is associated with red and orange forewing patterning in these butterflies. Nearby non-coding regions control its expression in *Heliconius* wing development [20] and these are the regions of divergence, while the coding region is a conserved homeobox gene. The forewings of *H. euphorbiae* are known for their pink flush in some regions of its distribution range (see moths in Fig. 3 of Hundsdoerfer, Lee et al. [3]), whereas red is not prominent in the grey wings of *H. vespertilio*.

The *WntA* signaling ligand is highly conserved at the amino acid level within *Heliconius* [20]) and is associated with the forewing black band. Variation is again more likely to be influenced by expression during wing development and is therefore found in the nearby control region [20].

The wing pattern gene *cortex* is associated with the yellow patterns on *Heliconius* wings [30] and has been suggested to underlie switches in color pattern across Lepidoptera [31]. For example, it has been shown that insertion of a transposable element (TE) into an intron of *cortex* results in industrial melanism [32], i.e. black morphs in the Peppered Moth (*Biston betularia*; Geometridae), corroborating its function across Lepidoptera, due to, e.g., the *cortex* TE insertion. This insertion [32], estimated to have occurred around 1819, could only manifest itself in the population through a selection pressure against the light-colored moths showing up on the dark trees and thus experiencing higher predation. In *Heliconius numata*, *cortex* lies within the *P*_1_ supergene inversion polymorphism and controls color pattern switching among mimicry morphs [26, 31].

Expression patterns of the two genes *distal-less* (Dll) and *engrailed* (En) [33] have been postulated to be a homologous developmental mechanism between butterfly and moth eyespots, also studied in saturniid moths of the genera *Antheraea* and *Saturnia* [19]. The gene *aristaless*, known to control wing color variation [34], has recently been shown to play an additional role in wing appendage formation during butterfly development [35], showing the seminal possibilities this field of research opens in understanding morphological evolution on the level of gene function. Overall, it is expected that a set of conserved, flexible wing patterning genes, possibly linked in a supergene, has also driven the rapid morphological diversification in the genus *Hyles*.

In contrast to *H. vespertilio* [12], in which the forewings have a naturally occurring knock-out appearance (as if the wing pattern genes were dysfunctional) of near-uniform grey wings lacking high-contrast patterning, *H. euphorbiae* shows important elements of the typical ground forewing pattern of the genus, which has been reconstructed as the ancestral set of characters for proto-*Hyles* [9]. By comparing the chromosome-level genomes of *H. euphorbiae* and *H. vespertilio*, the present work aims to provide reference genome data for future studies to understand the origin of *Hyles* wing patterns as phenotypic variability starting with several wing pattern control genes described above (e.g. [20, 23, 30]).

## Results

### *Hyles euphorbiae* genome assembly with chromosome-level scaffolding

The *H. euphorbiae* genome was assembled with Pac-Bio HiFi reads [36] combined with Hi-C data. Assembly contiguity statistics are summarized in Table 1. The final contig assembly consists of 56 scaffolds with an N50 of 18.2 Mb (321 contigs with an N50 of 2.76 Mb) and a size of 504 Mb and is available at NCBI (SRA accession number SRR17432892, BioProject PRJNA794104, BioSample SAMN24610150 and genome JALBCW000000000, JALBCX000000000, see data availability statement for itemized accession numbers). The annotated circular mitochondrial genome has a length of 15,322 bp. Final assembly statistics are summarized in Table 2.

**Table 1:**
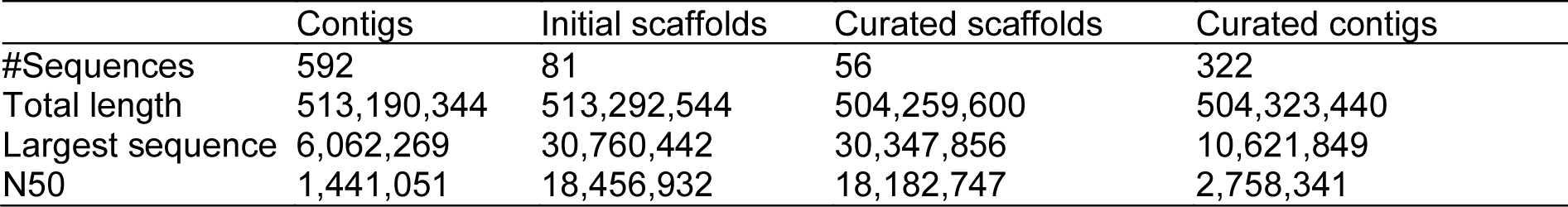
Contiguity statistics of *Hyles euphorbiae* genome.

|  | Contigs | Initial scaffolds | Curated scaffolds | Curated contigs |
| --- | --- | --- | --- | --- |
| #Sequences | 592 | 81 | 56 | 322 |
| Total length | 513,190,344 | 513,292,544 | 504,259,600 | 504,323,440 |
| Largest sequence | 6,062,269 | 30,760,442 | 30,347,856 | 10,621,849 |
| N50 | 1,441,051 | 18,456,932 | 18,182,747 | 2,758,341 |

**Table 2:** Assembly statistics of presented and related species. *Ms*: *Manduca sexta* [37], *Mt*: *Mimas tiliae* [14], *Lp*: *Laothoe populi* [15], *Hf*: *Hemaris fuciformis* [38], *Dp: Deilephila porcellus* [39], *He*: *H. euphorbiae, Hv*: *Hyles vespertilio*, the new assembly of *H. vespertilio* was used (based on PacBio sequence data from Pippel et al. [12]).

|  | *Ms* | *Mt* | *Lp* | *Hf* | *Dp* | *He* | *Hv* |  |
| --- | --- | --- | --- | --- | --- | --- | --- | --- |
| # scaffolds (>= 10000 bp) | 2822 | 30 | 33 | 34 | 31 | 47 | 380 |  |
| # contigs (>= 10000 bp) | 4977 | 41 | 135 | 74 | 36 | 313 | 526 |  |
| # scaffolds (>= 50000 bp) | 61 | 29 | 29 | 30 | 29 | 34 | 251 |  |
| # contigs (>= 50000 bp) | 1129 | 40 | 130 | 65 | 34 | 295 | 396 |  |
| Total length (>= 0 bp) Mb | 470.0 | 478.0 | 576.4 | 448.9 | 402.1 | 504.3 | 651.4 |  |
| Total length (>= 1000 bp) Mb | 470.0 | 478.0 | 576.4 | 448.9 | 402.1 | 504.3 | 651.4 |  |
| Total length (>= 5000 bp) Mb | 468.5 | 478.0 | 576.4 | 448.8 | 402.1 | 504.3 | 651.4 |  |
| Total length (>= 10000 bp) Mb | 463.1 | 478.0 | 576.4 | 448.8 | 402.1 | 504.3 | 651.4 |  |
| Total length (>= 50000 bp) Mb | 410.9 | 478.0 | 576.3 | 448.7 | 402.0 | 504.0 | 647.4 |  |
| Largest scaffold (Mb) | 21.11 | 25.98 | 29.55 | 25.57 | 22.66 | 30.35 | 37.22 |  |
| Largest contig (Mb) | 3.17 | 22.90 | 24.17 | 24.16 | 22.66 | 10.62 | 13.20 |  |
| Total length (Mb) | 470.0 | 478.0 | 576.4 | 448.9 | 402.1 | 504.3 | 651.4 |  |
| N50 scaffolds (Mb) |  | 14.25 | 17.90 | 21.13 | 16.55 | 15.07 | 18.18 | 22.14 |
| N50 contigs (Mb) | 0.425 | 17.80 | 6.97 | 10.68 | 14.88 | 2.76 | 7.26 |  |
| N75 (Mb) | 12.41 | 15.70 | 18.59 | 14.30 | 12.99 | 16.57 | 20.07 |  |
| L50 scaffolds (Mb) | 14 | 12 | 13 | 12 | 12 | 12 | 13 |  |
| L50 contigs (Mb) | 297 | 13 | 27 | 14 | 13 | 54 | 34 |  |
| L75 | 23 | 19 | 20 | 20 | 20 | 19 | 20 |  |
| # N's per 100 kb | 230.91 | 0.65 | 5.29 | 3.26 | 0.4 | 12.66 | 2.24 |  |
| BUSCO (lepidoptera_odb10): |  |  |  |  |  |  |  |  |
| complete, single-copy (%) | 91.8 | 98.5 | 98.5 | 98.4 | 98.6 | 97.9 | 95.4 |  |
| complete, duplicated (%) | 6.5 | 0.3 | 0.4 | 0.3 | 0.2 | 0.3 | 2.9 |  |
| fragmented (%) | 0.7 | 0.3 | 0.3 | 0.3 | 0.6 | 0.5 | 0.7 |  |
| missing (%) | 1.0 | 0.9 | 0.8 | 1.0 | 0.9 | 1.3 | 1.0 |  |

### Chromosome scaffolding with *Hyles vespertilio* Hi-C data

HiRise [40] scaffolding of *H. vespertilio* using more than 132 million read pairs (NCBI SRA accession number SRX14530528) yielded a scaffold N50 of 22.1 Mb. In total, 146 joins and only one break of the input assembly were introduced. The contact map clearly shows 29 well-supported scaffolds representing chromosomes (Fig. 2). Sizes of chromosomes of both *Hyles* species are of the same order of magnitude, but not identical (Table 3). The assembly of *H. vespertilio* is almost 147 Mb larger than that of *H. euphorbiae*.

**Fig. 2.**
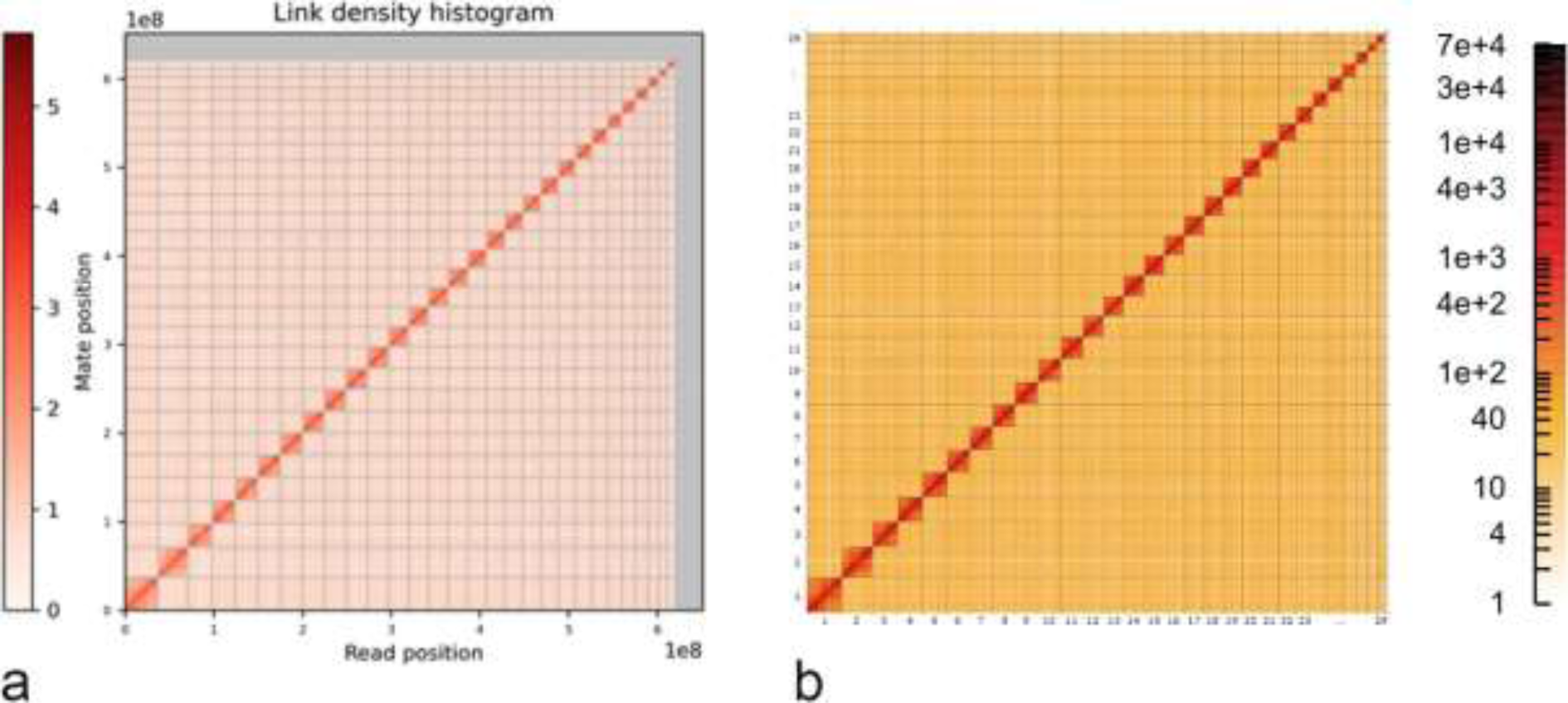
Contact map of a) *Hyles vespertilio* (Dovetail HiRise analysis) and b) *Hyles euphorbiae* (as described in M&M). Chromosome-level scaffolding clearly supports 29 scaffolds representing chromosomes. Chromosomes are ordered by size from bottom left (largest) to top right (smallest).

**Table 3:** Chromosome sizes of final assemblies and chromosome numbering in homology to *Bombyx mori*. Chromosome 1 corresponds to the Z chromosome [41]. Footnotes: * also contains parts of BmChr26, ** also contains parts of BmChr11, § also contains parts of BmChr23, # also contains part of BmChr24.

| *B. mori*<br>Chr.<br>number | *H. vesper-*<br>*tilio* Chr.<br>number | *H. vesper-*<br>*tilio* scaffold name<br>in browser | GenBank<br>Accession<br>number | Assembly<br>Length (bp)<br>incl. gaps | *H. eu-*<br>*phorbiae*<br>Chr.<br>Number | *H.*<br>*euphorbiae*<br>scaffold name<br>in browser | GenBank<br>Accession<br>number | Assembly<br>Length (bp)<br>incl. gaps |
| --- | --- | --- | --- | --- | --- | --- | --- | --- |
| BMChrZ | HvChrZ | ScWp86a_195_HRSCAF_280 | CM042833.1 | 37,218,281 | HeChrZ | sc_1 | CM041058.1 | 30,347,856 |
| BmChr2 | HvChr2* | ScWp86a_159_HRSCAF_226 | CM042805.1 | 24,553,817 | HeChr2 | sc_7 | CM041030.1 | 19,904,380 |
| BmChr3 | HvChr3 | ScWp86a_257_HRSCAF_395 | CM042806.1 | 20,065,448 | HeChr3 | sc_18 | CM041031.1 | 16,855,035 |
| BmChr4 | HvChr4 | ScWp86a_21_HRSCAF_27 | CM042807.1 | 25,450,662 | HeChr4 | sc_6 | CM041032.1 | 20,022,273 |
| BmChr5 | HvChr5 | ScWp86a_119_HRSCAF_165 | CM042808.1 | 26,040,715 | HeChr5 | sc_4 | CM041033.1 | 21,578,258 |
| BmChr6 | HvChr6 | ScWp86a_136_HRSCAF_194 | CM042809.1 | 22,136,963 | HeChr6 | sc_13 | CM041034.1 | 17,936,700 |
| BmChr7 | HvChr7 | ScWp86a_95_HRSCAF_134 | CM042810.1 | 17,182,831 | HeChr7 | sc_23 | CM041035.1 | 14,557,976 |
| BmChr8 | HvChr8 | ScWp86a_214_HRSCAF_310 | CM042811.1 | 21,873,546 | HeChr8 | sc_14 | CM041036.1 | 17,505,000 |
| BmChr9 | HvChr9 | ScWp86a_232_HRSCAF_355 | CM042812.1 | 24,188,504 | HeChr9 | sc_11 | CM041037.1 | 19,134,990 |
| BmChr10 | HvChr10 | ScWp86a_42_HRSCAF_61 | CM042813.1 | 24,738,224 | HeChr10 | sc_8 | CM041038.1 | 19,757,600 |
| BmChr11 | HvChr11 | ScWp86a_178_HRSCAF_255 | CM042814.1 | 22,098,621 | HeChr11 | sc_15 | CM041039.1 | 17,388,153 |
| BmChr12 | HvChr12 | ScWp86a_182_HRSCAF_262 | CM042815.1 | 26,852,440 | HeChr12 | sc_5 | CM041040.1 | 21,432,827 |
| BmChr13 | HvChr13 | ScWp86a_68_HRSCAF_94 | CM042816.1 | 24,145,169 | HeChr13 | sc_10 | CM041041.1 | 19,486,524 |
| BmChr14 | HvChr14 | ScWp86a_111_HRSCAF_155 | CM042817.1 | 20,079,433 | HeChr14 | sc_21 | CM041042.1 | 15,684,570 |
| BmChr15 | HvChr15 | ScWp86a_62_HRSCAF_86 | CM042818.1 | 26,014,704 | HeChr15 | sc_3 | CM041043.1 | 21,768,765 |
| BmChr16 | HvChr16 | ScWp86a_55_HRSCAF_78 | CM042819.1 | 18,679,767 | HeChr16 | sc_22 | CM041044.1 | 15,570,054 |
| BmChr17 | HvChr17 | ScWp86a_210_HRSCAF_302 | CM042820.1 | 24,352,496 | HeChr17 | sc_9 | CM041045.1 | 19,686,965 |
| BmChr18 | HvChr18 | ScWp86a_1_HRSCAF_1 | CM042821.1 | 20,909,092 | HeChr18 | sc_16 | CM041046.1 | 17,373,644 |
| BmChr19 | HvChr19 | ScWp86a_207_HRSCAF_299 | CM042822.1 | 20,353,206 | HeChr19 | sc_19 | CM041047.1 | 16,571,156 |
| BmChr20 | HvChr20 | ScWp86a_378_HRSCAF_523 | CM042823.1 | 16,497,537 | HeChr20 | sc_24 | CM041048.1 | 12,812,799 |
| BmChr21 | HvChr21 | ScWp86a_162_HRSCAF_230 | CM042824.1 | 21,114,399 | HeChr21 | sc_17 | CM041049.1 | 17,181,161 |
| BmChr22 | HvChr22# | ScWp86a_157_HRSCAF_223 | CM042825.1 | 33,800,775 | HeChr22 | sc_2 | CM041050.1 | 27,093,441 |
| BmChr23 | HvChr23 | ScWp86a_31_HRSCAF_41 | CM042826.1 | 22,933,977 | HeChr23 | sc_12 | CM041051.1 | 18,182,747 |
| BmChr24 | HvChr24 | ScWp86a_166_HRSCAF_234 | CM042827.1 | 11,894,872 | HeChr24 | sc_27 | CM041052.1 | 10,049,873 |
| BmChr25 | HvChr25 | ScWp86a_7_HRSCAF_7 | CM042828.1 | 19,247,757 | HeChr25 | sc_20 | CM041053.1 | 15,917,517 |
| BmChr26 | HvChr26** | ScWp86a_81_HRSCAF_111 | CM042829.1 | 9,224,630 | HeChr26 | sc_29 | CM041054.1 | 7,206,840 |
| BmChr27 | HvChr27 | ScWp86a_289_HRSCAF_430 | CM042830.1 | 14,304,874 | HeChr27 | sc_26 | CM041055.1 | 12,048,215 |
| BmChr28 | HvChr28# | ScWp86a_140_HRSCAF_198 | CM042831.1 | 15,210,208 | HeChr28 | sc_25 | CM041056.1 | 12,771,623 |
| n.a. | HvChr29§ | ScWp86a_137_HRSCAF_195 | CM042832.1 | 9,587,503 | HeChr29 | sc_28 | CM041057.1 | 7,786,824 |

### Chromosome structure in Sphingidae

Comparison of single copy orthologs in the *H. euphorbiae* genome assembly to those in *H. vespertilio* shows strong conservation of both synteny and chromosome structure, defined as conservation of gene order. This suggests that the last common ancestor of *Hyles* had 29 chromosomes that subsequently remained intact (Fig. 3a). These 29 chromosomes correspond to intact Merian elements, the ancestral linkage groups of Lepidoptera [42], with the exception of three simple fusion events between pairs of Merian elements (M17+M20, M30+M9, M24+M25). Next, the orthologs of *H. euphorbiae* were compared with those of *Deilephila porcellus* (Fig. 3b), both of which belong to Macroglossinae, a subfamily of Sphingidae. This also revealed a set of 29 orthologous chromosomes, suggesting that chromosome structure has remained stable throughout the evolution of this subfamily. To determine whether the stability of this set of 29 chromosomes extends throughout Sphingidae, we compared the orthologs of two further species, *Manduca sexta* (*n* = 28; Fig. 3c) of the subfamily Sphinginae and *Laothoe populi* (*n* = 28; Fig. 3d), which belongs to a third subfamily, Smerinthinae. This revealed that 27 chromosomes of *M. sexta* are orthologous and highly syntenic compared to *H. euphorbiae.* The remaining chromosome in *M. sexta* (CM026259.1) maps to two autosomes in *H. euphorbiae* (CM041057.1, CM041033.1), thus accounting for the difference in karyotype. This chromosome in *M. sexta* corresponds to a M17+20+M29 fusion. Similarly, one chromosome of *Laothoe populi* (HG992147.1) is orthologous to two autosomes in *H. euphorbiae* (CM041039.1, CM041057.1). This is due to a fusion between M29 and M4. Therefore, two independent fusion events have occurred in *L. populi* and *M. sexta*. This would suggest that the Sphingidae originally had a karyotype of *n* = 29 that has remained largely intact with a limited number of fusion events in extant species. Moreover, the strong conservation of gene order in *H. euphorbiae* compared with *M. sexta* illustrates limited intrachromosomal rearrangements since the last common ancestor of the Sphingidae.

**Fig. 3.**
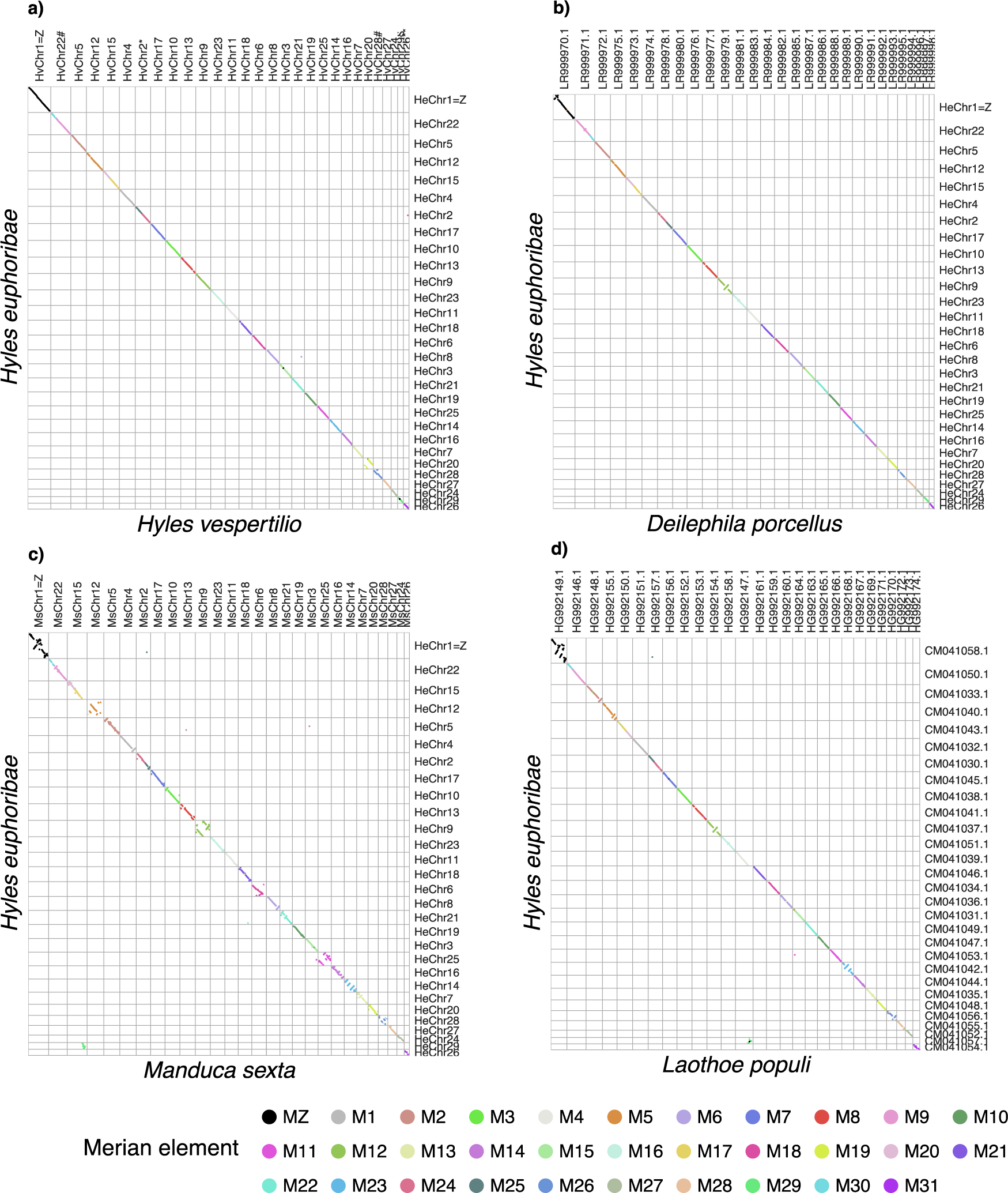
Oxford plots for a.) *Hyles vespertilio,* b.) *Deilephila porcellus* and c.) *Manduca sexta* d.) *Laothoe populi* versus *Hyles euphorbiae*. Comparison of assemblies demonstrate strong conservation of both linkage groups and gene order in Sphingidae. Orthologs are colored by Merian element identity. Chromosomes are ordered by decreasing size in *Hyles euphorbiae*.

### Genome alignments

The two species of the genus *Hyles* have one more chromosome than *B. mori* (*n* = 28), but the alignment between *H. vespertilio* and *B. mori* can still be illustrated (Fig. 4a) to allow comparison by visual inspection. Chromosome proportion values of this alignment are reported in Table S1.

**Fig. 4.**
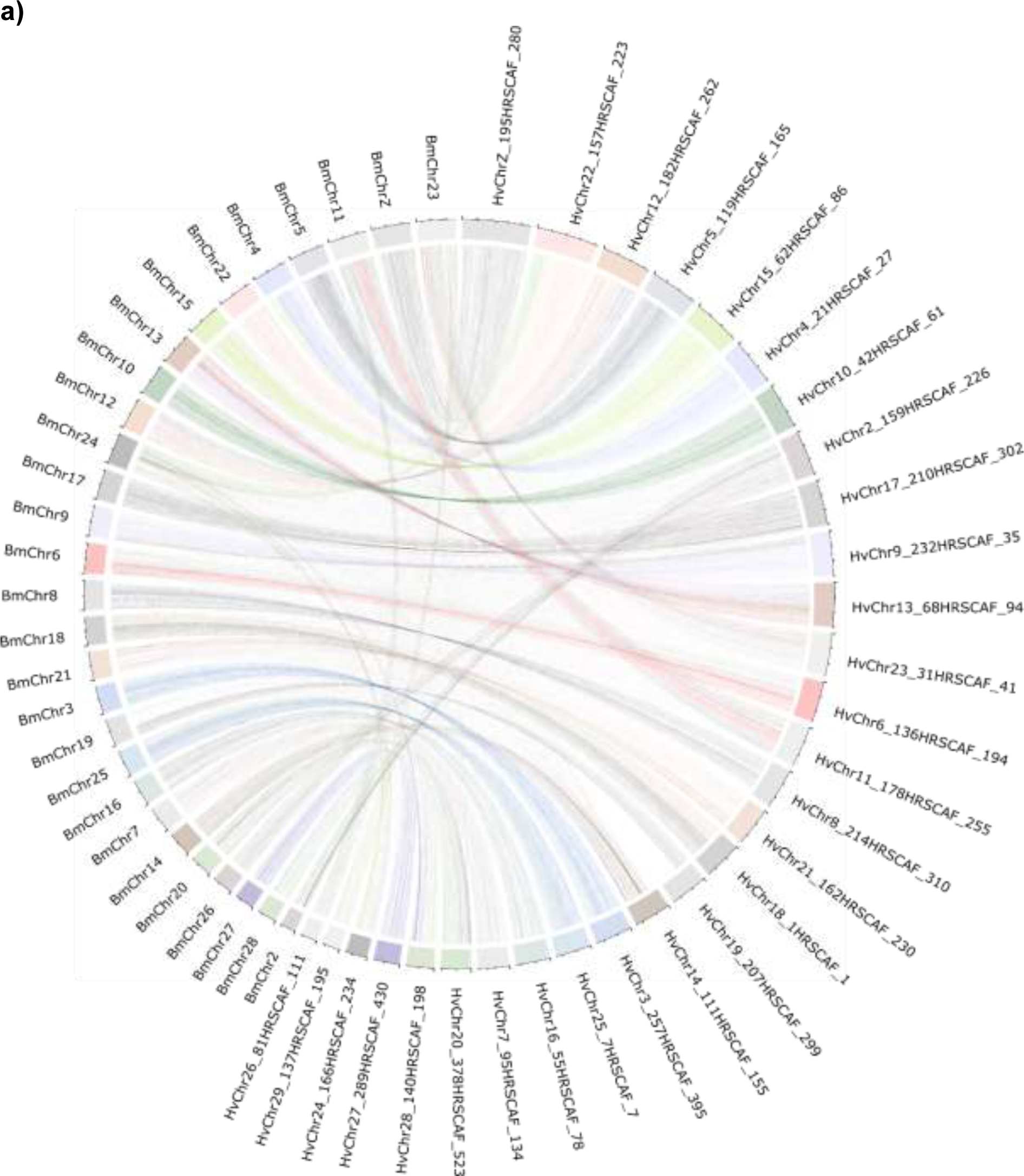
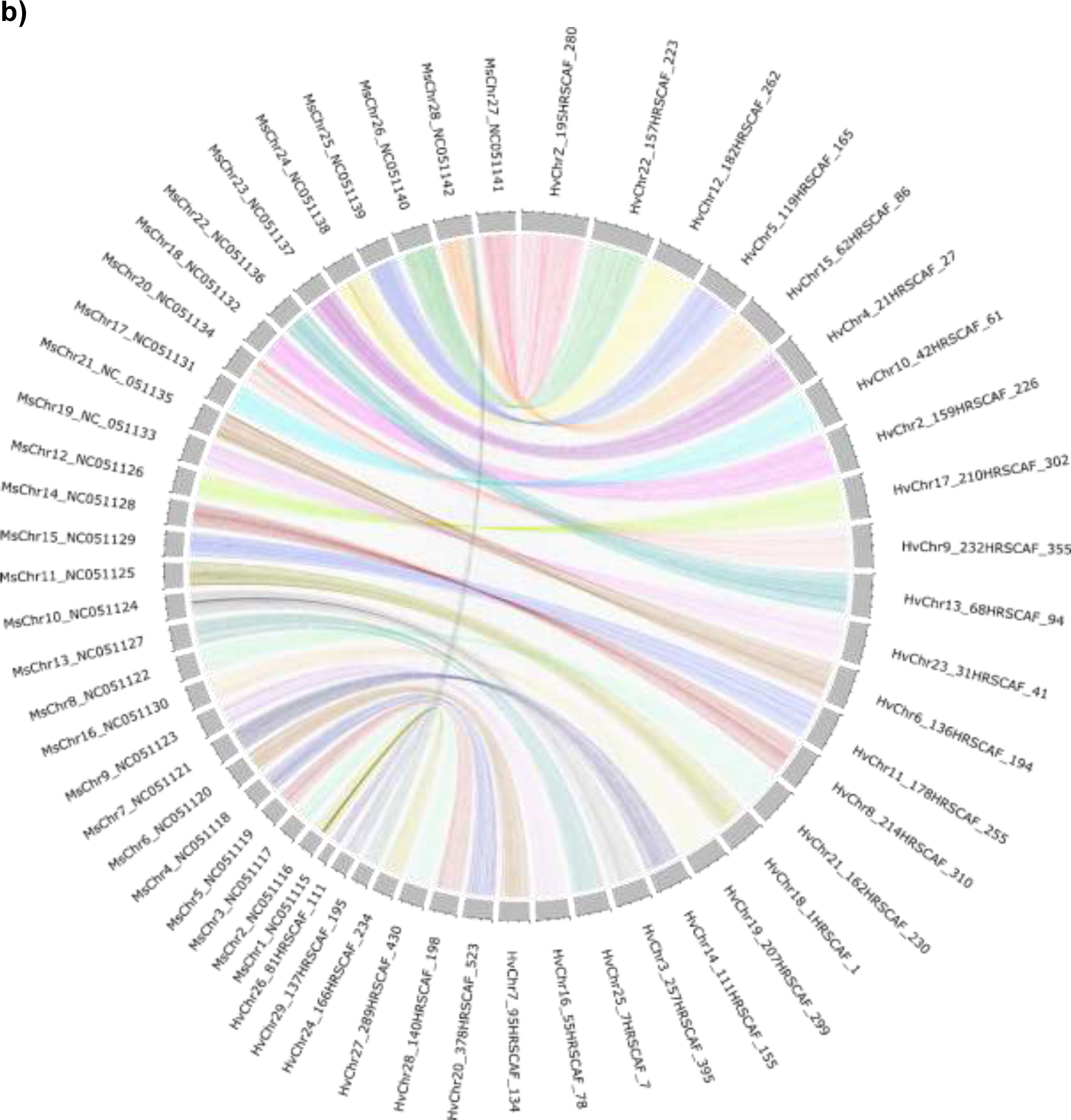
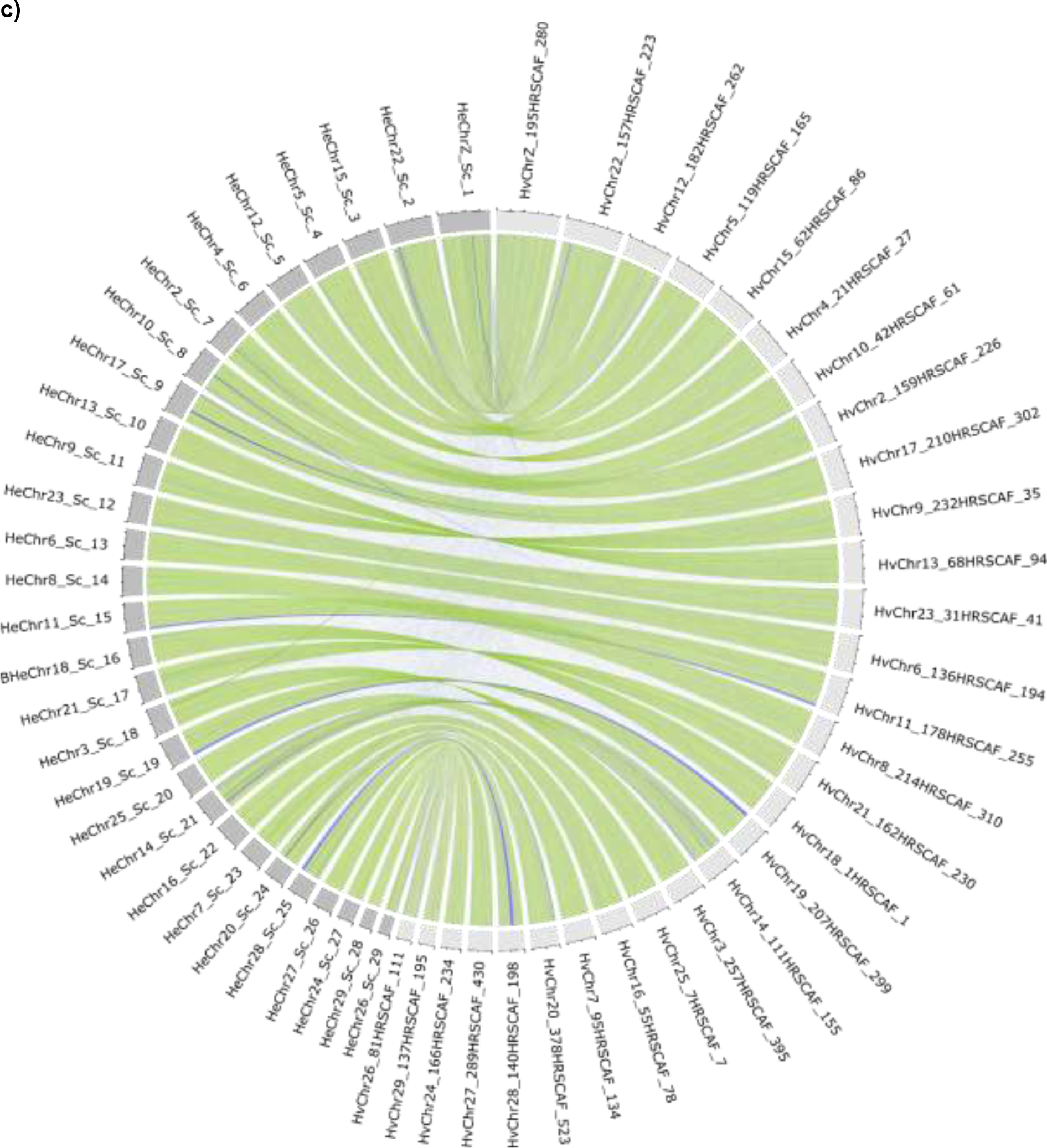
CIRCOS plots of genome alignments at the chromosome level. The Z chromosome corresponds to chromosome 1. Chromosomes of *H. vespertilio* are ordered by size. The letters “ScWp86a” in the scaffold names of *Hyles vespertilio* are omitted for clarity. **a**) *Bombyx mori* and *H. vespertilio* with automatic coloring using shinyCircos to facilitate correlation. **b**) *Manduca sexta* and *H. vespertilio* with coloring by chromosome. **c**) *H. euphorbiae* and *H. vespertilio*, with the 29 chromosomes of each species (*H. vespertilio* light grey boxes in the outer rim, *H. euphorbiae* dark grey boxes). The color of the links represents the strand orientation of the chromosomes of *H. euphorbiae* compared to the *H. vespertilio* strand orientation (green means +; blue means -).

CIRCOS plots of the alignment between the two species show well-preserved synteny, except for some chromosomes that could be involved in chromosome rearrangements. Chromosome 2 (HvChr2) of *H. vespertilio* corresponds to chromosomes 2 and 26 of *B. mori* (BmChr2, BmChr26). Chromosome 24 of *B. mori*, BmChr24, corresponds to parts of *H. vespertilio* chromosomes 22, 24, and 28 (HvChr22, HvChr24, HvChr28), and chromosome 11 of *B. mori* (BmChr11) corresponds to parts of *H. vespertilio* chromosomes 11 and 26 (HvChr11, HvChr26). These chromosome rearrangements between *B. mori* and *H. vespertilio* are consistent with those between *B. mori* and *M. sexta* [37, 43], corroborating the strong synteny between the two hawkmoth species, *H. vespertilio* and *M. sexta* (Fig. 4b). Chromosome 23 of *B. mori* (BmChr23) is split into *H. vespertilio* chromosomes 23 and 29 (HvChr23, HvChr29), resulting in an additional chromosome in *H. vespertilio* (HvChr29). In *M. sexta*, which has 28 chromosomes, HvChr29 and HvChr15 correspond to MsChr28, thus showing a different combination.

The plot comparing the 29 chromosome sequences of *H. vespertilio* and *H. euphorbiae* (Fig. 4c) illustrates the highly conserved synteny within the genus *Hyles* in the definition of collinearity, the conservation of block order in the two sets of chromosomes. The larger size of the *H. vespertilio* genome based on the genome size estimation and its longer assembly than that of *H. euphorbiae* are reflected by the larger size of every chromosome (Fig. 4c).

### Annotation and methylation profile

The proportion of the genome covered by the major classes of repetitive elements is 37% and 47% for *H. euphorbiae* and *H. vespertilio*, respectively (Table 4, Fig. 5).

**Table 4.** Assembly lengths and proportion of repeats of the available (on 14.10.2022) genomes in comparison (a new assembly of *H. vespertilio* was used, based on PacBio sequence data from Pippel et al. [12]).

| Species | Total length | Masked [%] |
| --- | --- | --- |
| *Manduca sexta* [37] | 470,036,997 | 27.8 |
| *Mimas tiliae* [14] | 472,718,480 | 46.7 |
| *Laothoe populi* [14] | 549,962,493 | 47.7 |
| *Hemaris fuciformis* [38] | 441,141,813 | 33.2 |
| *Deilephila porcellus* [39] | 389,730,088 | 27.5 |
| *H. euphorbiae* | 504,310,614 | 37.1 |
| *H. vespertilio* | 651,427,907 | 46.6 |

**Fig. 5.**
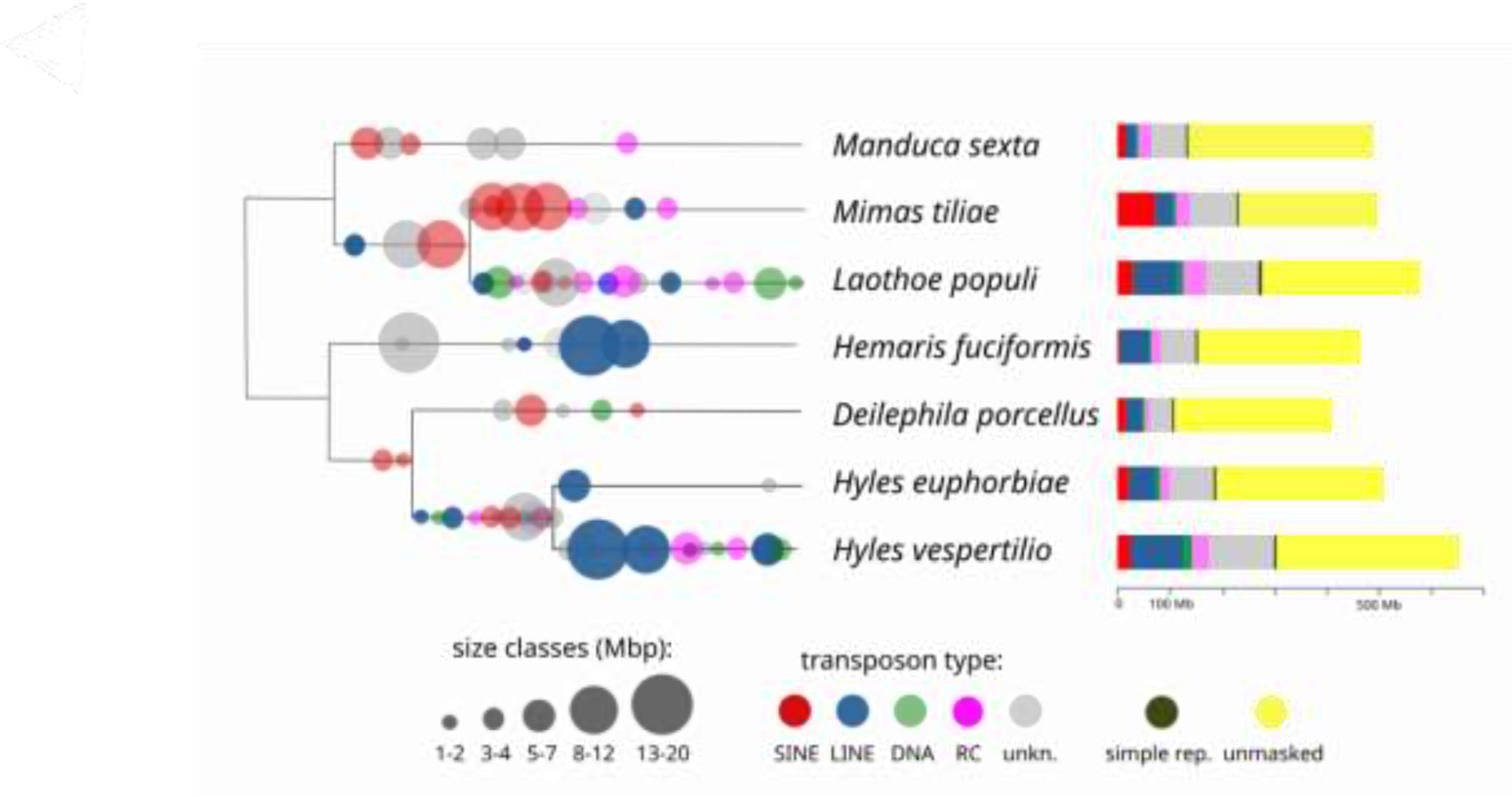
Transposon bursts mapped on a phylogenetic tree drawn by hand according to backbone topology in [44] in relative order (due to sequence diversity) and proportional repeat contents of seven sphingid species with available genome data (transposon types: SINE: short interspersed elements, LINE: long interspersed elements, DNA: DNA transposons, RC/Hel: rolling-circle transposons, e.g. helitrons).

Thus, *H. vespertilio* has a much larger genome (651 Mbp to 504 Mbp in *H. euphorbiae*) and a higher proportion of repetitive DNA than *H. euphorbiae*. A comprehensive comparison of transposon content in seven species of the family Sphingidae (Fig. 5) demonstrates the dynamics of repetitive elements and the contribution of recent bursts in transposable element families to genome size expansion in *H. vespertilio* and independently in *Laothoe populi*.

The number of genes per chromosome is very similar in the two species’ genomes presented here (Fig. 6), but exhibits a wide range from a minimum of 169 genes in both species on chromosome 29 to 826 / 798 genes in *H. vespertilio* / *H. euphorbiae* on chromosome 22.

**Fig. 6.**
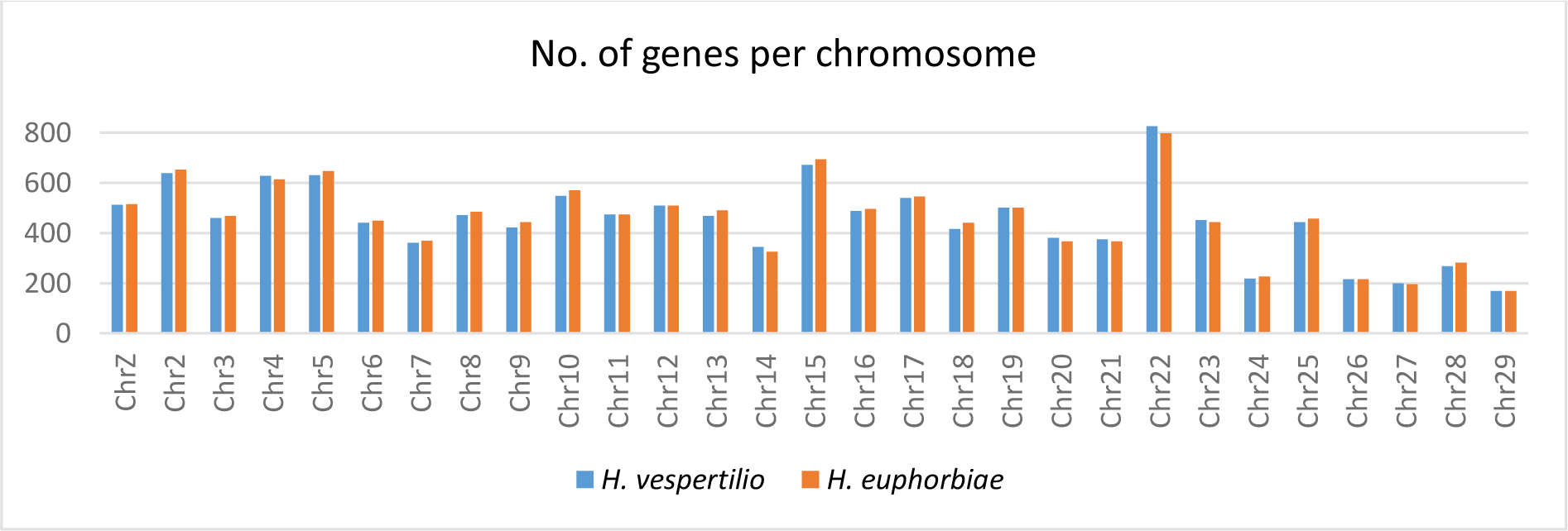
Number of genes per chromosome for *H. euphorbiae* and *H. vespertilio* in comparison.

The mitochondrial genome of *H. euphorbiae* which was assembled with the mitoHifi pipeline [45, 46] based on the closely related species, *Theretra oldenlandiae*, is illustrated in Fig. S1. It was annotated using MITOS and contains 13 protein coding genes, two ribosomal RNA (rRNA) genes plus 22 transfer RNA (tRNA) sequences and the control region.

The methylation profile of *H. euphorbiae* (track “HL Methylation” in the genome browser; e.g. Fig. S5a) shows modification probabilities for every site. Base modifications with probability of modification < 50% (hypomethylation) are colored blue and base positions that have a high probability to be methylated and thus not be expressed are marked red (base modification > 50% probability, hypermethylation).

### Wing pattern genes

The wing pattern genes *optix, WntA*, *cortex, aristaless* and *distal-less* (accession numbers of reference sequences are presented in Table S2) were identified with high confidence using BLAT [47, 48]; only one hit was found for each with thresholds of identity above 85% and bit score above 420.

The hits of the *Heliconius erato optix* protein sequence revealed a position on chromosome 23 in *H. euphorbiae* (HeChr23, “sc_12: 1,582,862-1,583,680”, size: 819 bp) and *H. vespertilio* (HvChr23, “ScWp86a_31_HRSCAF_41: 20,799,551-20,800,375”, size: 825 bp). No repetitive elements were annotated within the area of the coding sequence in either species (Fig. S2). However, upon zooming out 10x or 100x a high amount of indels are seen in the genome browser (400 KB up-& downstream of the coding sequence, Fig. S3).

The *Heliconius himera WntA* amino acid sequence (Table S2) mapped to the *Hyles* genomes returned a position on chromosome 28 in *H. euphorbiae* (HeChr28, “sc_25: 5,829,059-5,837,439”, spanning six exons and 8,381 bp; Fig. S4a; annotated by transcript LOC115447391_rna-XM037443222.1.47 of *Manduca sexta* spanning seven exons and 17,247 bp) and *H. vespertilio* (HvChr28, “ScWp86a_140_HRSCAF_198: 8,200,403-8,210,485”, spanning six exons and 10,083 base pairs, Fig. S4b; annotated by evm07703.91 spanning seven exons and 14,395 bp). Views of the *WntA* BLAT results are illustrated in Fig. S4. There are numerous differences in the repeats between the two *Hyles* species. For example, *H. vespertilio* contains a 443 bp long DNA repeat insertion (“ScWp86a_140_HRSCAF_198: 8,200,940-8,201,382”; rnd-4_family-526; family TcMar-Tc1, pos. 512-1311 in the repeat) in intron 2 that *H. euphorbiae* lacks. The genomic interval spanning the exons and introns of *WntA* is inverted in *H. euphorbiae* (or *H. vespertilio*, depending on the polarity definition).

BLAT search of the *M. sexta cortex* protein sequence (8 exons) resulted in a match corresponding to 7 exons on chromosome 17 in both species [Fig. S5; *H. euphorbiae* (HeChr17, “sc_9: 7,671,034-7,674,090”) and *H. vespertilio* (HvChr17, “ScWp86a_210_HRSCAF_302: 14,798,018-14,801,349”)]. Views of the *cortex* BLAT results are illustrated in Fig. S5. The methylation profile in *H. euphorbiae* (Fig. S5a) indicates normal transcription activity in this stretch of exons, as the probabilities of base modifications are all < 50% (blue). The position, quality, and quantity of repeats in introns differ between the two species. For example, the first intron in *H. euphorbiae* contains no repeats, whereas in *H. vespertilio* it contains two LINEs in addition to a 211 bp stretch (ScWp86a_210_HRSCAF_302: 14,798,399-14,798,609; Fig. S5b) of unknown repeats (named rnd-1_family-40). It overlaps downstream with the LINE of family L2, which shows an irregular simple repeat of 5’ TATT. It is in intron 1 of the *cortex* X1 sequence of *M. sexta* (annotated with *M. sexta* transcript LOC115451467_rna-XM_030179809.2.8). Again, the area surrounding the gene *cortex* is inverted in *H. euphorbiae* (or *H. vespertilio*, depending on the polarity definition). The 100 kb views of the *cortex* BLAT results are illustrated in Fig. S6 and reveal a high number of repeats in the vicinity of the stretch of gene exons in both species.

The BLAT search of *M. sexta aristaless* (in 4 variants, Table 2) revealed the gene to occur on chromosome 4 in both species (HeChr4, “sc_6: 4,437,062-4,474,302”, spanning 37,241 bp) and *H. vespertilio* (HvChr4, “ScWp86a_21_HRSCAF_27: 5,766,997-5,884,695”, 5 exons within a large stretch of 117,699 bp). The first intron of *aristaless* contains a repeat motif in *H. euphorbiae* that is not found in *H. vespertilio* (Fig. S7). Apart from this difference, the positions of the other repeats within the stretch of exons are very similar in both species, but the stretches of repeats are longer in *H. vespertilio* (scale of 2 kb in both views of Fig. S7). Upon zooming out, the large diversity in repeats in the vicinity of this gene becomes apparent (Fig. S8), as well as the large stretches of non-coding DNA between the numerous exons.

The BLAT result of *distal-less* (Table 2) reveals a position on chromosome 2 in both species (HeChr2, “sc_7: 16,368,465-16,384,630”, 2 exons within 16,166 bp) and *H. vespertilio* (HvChr2, “ScWp86a_159_HRSCAF_226: 4,646,872-4,661,762”, 2 exons within 14,891 bp). The single intron between the two exons contains repeats in both species, however very different types in different positions and extents (Fig. S9).

### Karyotype

Analysis of male mitotic chromosomes stained by FISH with a telomeric probe (telomere-FISH) showed that the karyotype of *H. euphorbiae* is composed of 2*n* = 58 chromosomes (Fig. 7a). As typical for Lepidoptera, the chromosomes are of the holokinetic type, i.e. they lack a primary constriction (centromere) and are morphologically uniform, differing only in size. Chromosome number was confirmed by analysis of meiotic nuclei at the pachytene stage, where homologous chromosomes pair and form elongated bivalents. Pachytene complements, stained by GISH in combination with telomere-FISH, showed a haploid number of 29 bivalents in both sexes (Fig. 7b, c). In addition, GISH identified a WZ sex chromosome bivalent in pachytene oocytes by labeling a large portion of the W chromosome with the female genomic DNA (gDNA) probe (Fig. 7c), whereas no bivalent was identified in pachytene spermatocytes (Fig. 7b). These results clearly show that *H. euphorbiae* has a WZ/ZZ (female/male) sex chromosome system, which is common in Lepidoptera. It should be noted that the WZ bivalent is relatively long (Fig. 7c), suggesting that the W and Z chromosomes are among the largest chromosomes in the *H. euphorbiae* karyotype.

**Fig. 7.**
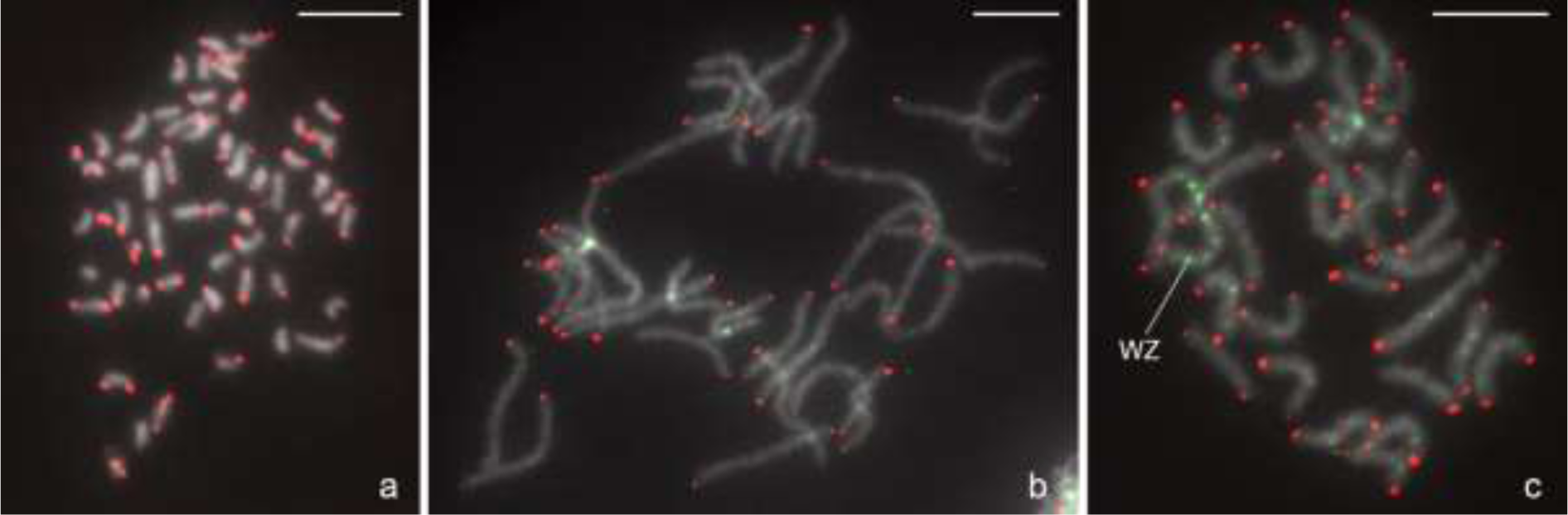
Molecular cytogenetic analysis of *Hyles euphorbiae* chromosomes. Hybridization signals of the Cy3-labeled (TTAGG)_n_ telomeric probe (red) indicate chromosomal ends (**a**–**c**), and the fluorescein-labeled female gDNA probe (green) identifies the sex chromosome system (**b** and **c**). Chromosomes were stained with DAPI (grey). (**a**) Male mitotic prometaphase stained with telomere-FISH showing a diploid chromosome number of 2*n* = 58. (**b**) Male pachytene complement stained with a combination of GISH and telomere-FISH showing 29 bivalents, but without any bivalent highlighted, indicating a ZZ sex chromosome constitution. (**c**) Female pachytene complement stained with a combination of GISH and telomere-FISH, showing 29 bivalents, including the WZ sex chromosome pair identified by the W chromosome highlighted with the female gDNA probe. Bar = 10 µm.

### Chromosome size estimation from karyotype image data

Chromosome size estimation is based on Fig. 7c, bivalents from a female gonad cell at the pachytene stage. The chromosome size estimates are shown in Fig. 8. They corroborate the WZ bivalent as the largest chromosome. Based on semi-automated image processing, the software package napari-karyotype [49] relies on threshold-based image segmentation to detect chromosome-related components. Identified chromosomal objects are surrounded by red rectangles and labeled with the estimates.

**Fig. 8.**
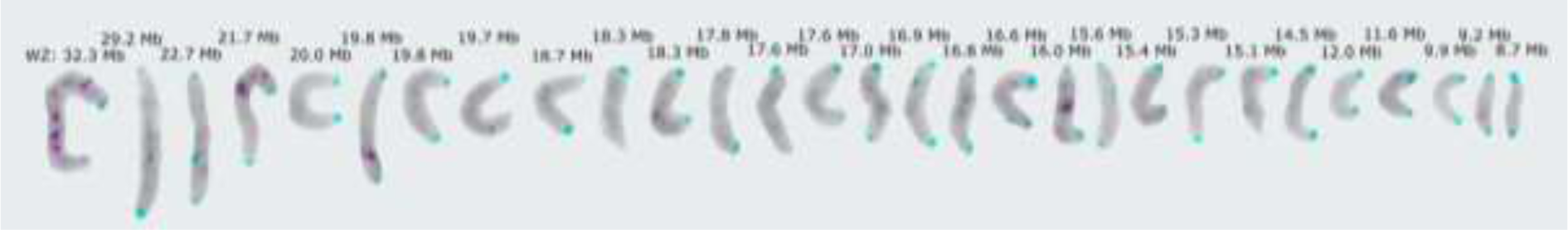
Annotated chromosome bivalents from a female pachytene image of *Hyles euphorbiae* using the software package napari-karyotype. Chromosome bivalents were aligned manually according to the estimated size. The largest chromosome pair represents the sex chromosomes, in this case a WZ bivalent. Blue and red dots along the chromosomes correspond to the location of the telomeric probe and female gDNA probe, respectively, from Fig. 7.

### Genome size estimation

Genome sizes of *H. vespertilio* and *H. euphorbiae* were measured using flow cytometry in three replicates, each from a single individual. The average results of 0.575 pg and 0.484 pg showed that the 1C DNA values were 562 Mb and 472 Mb, respectively (1 pg = 978 Mb).

## Discussion

### Assembly quality

The chromosomal genome assemblies of *H. euphorbiae* and *H. vespertilio* presented here are highly contiguous, as demonstrated by the scaffold N50 values exceeding those of the sequenced close relatives *M. sexta* [37] and *B. mori* [41]. The percentage of assembly length that could be placed in chromosome scaffolds is highest in *Hyles euphorbiae* (Table 2). Both metrics underline the high quality of the two new assemblies for the *Hyles* genus. Gene content is similarly high in all compared assemblies.

### Comparison of the two new *Hyles* genomes

The large difference in genome size estimates between the two species within the genus *Hyles*, i.e. *H. vespertilio* with 562 Mb being ∼20% larger than that of *H. euphorbiae* with 472 Mb in the flow cytometry measurements, was unexpected due to the short divergence time between the two species [50]. *Hyles lineata* is one of the first *Hyles* species to diverge within the genus [5] and has an even smaller flow cytometry genome size of 450 Mb (0.46 pg) [51]. The potentially smaller genome size of this third *Hyles* species suggests that the larger genome of *H. vespertilio* is the result of a recent proliferation of repetitive elements and is a derived character.

Although the values of genome sizes based on assembly lengths are not directly comparable to genome size values based on flow cytometry due to technical differences in methodology, the relative sizes are of the same order of magnitude. The assembly length of *H. vespertilio* with 651 Mb is ∼30% longer than that of *H. euphorbiae* with 504 MB, although the number of genes is very similar in both species (Fig. 6).

The genome of *H. euphorbiae* presented in this work has many fewer repetitive elements, in particular LINEs, than *H. vespertilio*. The high repeat content in all four species compared, *H. euphorbiae, H. vespertilio, M. sexta*, and *B. mori,* is typical for lepidopteran genomes [52, 53]. However, the number of repeats varies between species. Previous research has shown a correlation between the repetitive fraction of the genome and genome size within and between species [54, 55]. Indeed, the genome of *H. vespertilio* is the largest in our comparison and the genome with the most extensive repeat content, whereas the other sphingid genomes show lower numbers of repeats, correlating with their smaller genome size. Therefore, as described in previous research, the repetitive elements found here are thus likely the driving force for the expansion of genome size, possibly due to a positive feedback loop that allows these elements to spread more easily in large genomes [56]. In addition, the repeatome is thought to play a significant role in genetic innovation and speciation [57–61]. This could explain why *H. vespertilio* is known to be one of the most isolated species in the genus of hawkmoths with respect to the lack of hybridization with numerous other *Hyles* species. In contrast, *H. euphorbiae* is well known for its wide distribution and frequent hybridization with even distantly related species [2, 50, 62].

Since the two species are separated by a *p*-distance of only 2.1% in the neutral marker genes *COI* and *COII* [50], corresponding to a split timing of only 4.2-5.6 Mya (according to the respective molecular clocks of 1.1–1.2% per million years per lineage, resulting in 2.3% sequence divergence between species pairs per million years of separation [63]), high synteny of genes between *H. vespertilio* and *H. euphorbiae* in the nuclear chromosomes set was expected (Figs. 3, 4). Indeed, the illustrated genome alignments (Figs. 3, 4) show how similar the two *Hyles* genomes are. The mitochondrial genome of *H. euphorbiae* (Fig. S1) is also very similar to that of *H. vespertilio* (see [13]). Comparison of single copy orthologs in each genome demonstrates conservation of 29 orthologous chromosomes. These chromosomes closely correspond to the 32 ancestral chromosomes of Lepidoptera (Merian elements)[42], with the exception of three fusions, thus accounting for the reduction in chromosome number. Moreover, we observe a high degree of synteny, defined as conservation of gene order within each chromosome, in line with other studies[42, 52, 64, 65]. Extending the comparisons beyond *Hyles* suggests that these 29 chromosomes have remained stable throughout Sphingidae, with the exception of two independent fusion events in *M. sexta* and *L. populi,* which are absent in *Hyles* and *D. porcellus.* Such strong conservation of chromosome structure has been previously observed in Nymphalidae [66]. Our analyses presented here suggest that the Sphingidae chromosomes are also remarkably syntenic over long periods of evolution.

### Karyotype, chromosome taxonomy

In addition to examining conservation of chromosome structure through comparison of single copy orthologs to Merian elements in each genome, we also performed whole genome alignments. Whole genome alignments between each *Hyles* genome and that of the well-studied species *B. mori* were done using positional information of synteny blocks (Table S1), in order to assign homology between *B. mori* and *Hyles* chromosomes, enabling chromosome naming based on *B. mori*. This resulted in the robust chromosome taxonomy based on *B. mori* (presented for the two *Hyles* species studied in Fig. 4c).

Given the young age of the genus (the oldest intra-genus split is 10.8-12.4 Mya, based on *p*-distance [2]), it can be assumed that the chromosome taxonomy also applies to the entire genus *Hyles*. We propose our protocol of homology inference based on genome alignments as the gold standard for chromosome taxonomy. The most recent hawkmoth assemblies [14, 39], including that of *M. sexta* [37], use only arbitrary numbers according to chromosome size. A table showing homology of chromosomes, accession and scaffold numbers is provided in Table S3. Using this comparative approach as a standard in bombycoid chromosome taxonomy will greatly facilitate discussions based on genomic comparisons for specific research questions, such as the homologies of wing pattern genes in the present study. In early genetic studies, three chromosomes of *B. mori* (BmChr11, BmChr23 and BmChr24) were often interpreted to be split in other species, increasing the chromosome number compared to *B. mori*, e.g. [43, 67, 68]. The recent discovery of 32 Merian elements, the ancestral linkage groups of Lepidoptera, provides a framework within which to understand the chromosomal changes in *B. mori* relative to other species. The ancestral state of Ditrysia is *n*=31 due to an ancient fusion between two Merian elements. Therefore, *B. mori* has a reduced chromosome number relative to other Ditrysians due to a subsequent three fusions.

The whole genome alignments between *B. mori* and *Hyles* demonstrate that these two groups have undergone independent sets of fusions, leading to the reduced chromosome numbers. This is consistent with the previous observationthat three *B. mori* chromosomes are often split in other species.

The *H. euphorbiae* chromosome images obtained had a sufficient clarity to be annotated with a size estimate by napari-karyotype (Tab. 3) [49]. However, it should be noted that this was more difficult than expected because the chromosome bivalents at the pachytene stage were touching each other in the image and thus had to be extracted manually. Without this step, the assignment of object vs. background would have been inaccurate. Furthermore, chromosomes are flexible structures and their length depends on the stage, the degree of condensation and also on the preparation methods. Therefore, their measured length does not always correspond to their size, which is especially true for meiotic chromosomes at the pachytene stage [69]. In order to annotate the chromosome images of the karyotype with the chromosome numbers of the *B. mori* chromosome taxonomy, it will be necessary to implement further in-situ-hybridization with gene-specific fluorochrome-labeled probes following Yasukochi et al. [43] (their Figure 2) in the future.

### Wing pattern genes

The location of numerous wing pattern genes has been well studied in Nymphalidae. Here, we infer the likely locations of *optix*, *WntA*, *cortex, aristaless* and *distal-less* in *Hyles* based on amino acid or the homologous *M. sexta* nucleotide similarity and characterize their genomic context.

#### optix

*optix* and surrounding genes are highly conserved in Lepidoptera [70]. Zhang et al. [28] have shown that *optix* knock-outs exhibit complete replacement of color pigments by melanins, resulting in black and grey butterflies. In contrast, the high percentage of indels found in the alignment up-and downstream of *optix* between the genomes of *H. vespertilio* and *H. euphorbiae* may support the previously proposed hypothesis [71] that wing patterns are indeed controlled by cis-regulatory elements near the position of *optix*. As more sequence data from more individuals become available, *F_ST_* plots can be calculated to corroborate correlations between particular candidate regions. Although *Manduca sexta* is the reference moth genome most apt in the comparative framework presented here, the wing pattern of this species also consists of black and grey, without colored stripes or spots. Thus, the inversion of *optix* on chromosome 23 in *H. euphorbiae* with respect to *H. vespertilio* and *M. sexta*, could possibly also be the other way round, in that *H. euphorbiae* with its cream, olive-brown forewings, sometimes with pink, holds the gene with normal function, and *H. vespertilio* and *M. sexta* have a non-functional inversion. The hypothesis if *optix* is part of a supergene will be studied in more detail by targeted resequencing of phenotypes in comparison. However, the causal relationship between grey wings in *H. vespertilio* and *optix* and surrounding gene regions could only be answered by analyzing knock-outs generated using the CRISPR/Cas9 system.

#### WntA

The highly varying amounts, classes, and families of repeats between the two *Hyles* species sequenced suggest that the function of this gene could have been corrupted in the grey *H. vespertilio*, as this species has the larger genome and more repeats. Sequence data from more individuals are needed to pinpoint a candidate region and establish a robust correlation.

#### cortex

Another striking difference between the two species near the stretch of 8 *cortex* exons is the insertion of 229 bp, which was marked as an unknown repeat (Fig. S5b) on chromosome 17 in *H. vespertlilio*. Mapping this fragment to the genome of *H. euphorbiae* using BLAT yielded 200 hits between 89.8-96.7% identity on each chromosome and 203 hits on the new genome of *H. vespertilio* (data not shown), indicating the wide distribution of this repeat region across the genomes. It was also found in the very closely related macroglossine species *Deilephila porcellus* [39] and *Hemaris fuciformis* [38], as well as several hits in the genome of the smerinthine *Mimas tiliae* [14], a phylogenetically somewhat more distant species. Again, *cortex* is found to lie on an inversion in *H. euphorbiae* (although in this comparative framework, it is possibly the other way round), directing thoughts to a possible connection within a second supergene (since it is on a different chromosome than *optix*). The two new *Hyles* reference genomes open doors to investigating these possibilities. Further analyses and comparisons between the genomes presented here and others becoming available are expected to lead to a better understanding of the mechanisms controlling wing patterns in the family Sphingidae.

#### aristaless & distal-less

The two genes appear similarly diverse between the species with respect to the number and type of repeats. The two *Hyles* moths studied do not show eyespots in their wings, nor do other species of the genus. However, further studies could clarify whether these genes control the simple brown spots on the forewings in *H. euphorbiae* and many other species of this genus by understanding if the genes are not expressed or possibly silenced in *H. vespertilio*.

## Conclusions

Previous studies had already shown that similarity of wing patterns does not correlate with phylogenetic relationships in the genus *Hyles* [1, 2, 5], neither as currently defined by morphology nor as reflected by molecular phylogeny. Morphologists have long argued that they must rely on phenotypic characters in the absence of molecular data, and have often included striking differences in wing patterns in their species descriptions. In *Hyles*, however, the evolution of wing pattern characters does not reflect the evolution of the species. Of course, gene trees are not species trees [72], so even traditional genetic analyses, e.g. [1, 2, 5], do not necessarily reflect the true tree.

In this study, we present two high-quality annotated chromosome-level assemblies and report the presence, sequence, and location of wing pattern genes. Using reference genomes of *H. euphorbiae* and *H. vespertilio*, which have very different forewing patterns and coloration, we open up possibilities for studying the evolution of wing patterns based on a numerically evaluable, objective data source. The genes *optix* (on chromosome 23), *WntA* (on chromosome 28), *cortex* (on chromosome 17), *aristaless* (on chromosome 4) and *distal-less* (on chromosome 2) promise to be of interest in a comparative study of numerous *Hyles* individuals, as the genomic regions surrounding these genes show high numbers of indels potentially corrupting their function and inversions were detected between the two *Hyles* species. Since the five genes compared in this study all lie on different chromosomes, much work is still needed until we understand the basis of the changes between e.g. the two prominent *Hyles* forewing patterns described in the introduction, let alone the additional often unique forewing patterns in other species. The results of our comparative repeatome analysis confirm that repetitive elements are likely the driving force for the expansion of genome size in *Hyles vespertilio* compared to *H. euphorbiae*. The chromosome-level genomic data provided in this study for these two species provide reliable references in the family Sphingidae for future studies involving as many species as possible to elucidate the evolution of forewing patterns in this group of Lepidoptera.

We strongly suggest the application of a standardized chromosome naming process for every newly sequenced genome to facilitate comparability and propose using a two-fold comparative genomics approach. The combination of CIRCOS plots based on pairwise alignments in a given system (such as *B. mori* for Bombycoidea) for chromosome taxonomy with Oxford plots based on Merian elements to infer and visualize directionality of chromosomal rearrangements, enables more precise evolutionary deductions.

## Material and Methods

### Insects

For karyotype analysis, specimens of the *H. euphorbiae* population from Greece (leg. P. Mazzei, Serifos) were reared in the laboratory (summer 2019). Several larvae and young pupae were used to make chromosome preparations (see below). For genome sequencing, one male specimen of *H. euphorbiae* (Fig. 1a) was collected near Berbisdorf (Germany) on 27 July 2021 and received the tissue voucher number MTD-TW-13387 in the Molecular Laboratory of the Senckenberg Naturhistorische Sammlungen Dresden, Museum of Zoology. A second, similar male moth from the same locality was placed as a voucher specimen in the main SNSD Entomology collection. The Hi-C data for *H. vespertilio* (Fig. 1b) was obtained from frozen (-80°C) tissue of the same individual from Vallonina, Italy (collected in 2018) that was used for the original genome publication in Pippel et al.[12]. The tissue was moved from the Max Planck Institute of Molecular Cell Biology and Genetics (Dresden), where it had the accession number LG2117, to the SNSD-MTD tissue collection, and received the voucher number MTD-TW-13386.

### Karyotype analysis

Spread chromosome preparations were made as described by Yoshido et al. [73]. Mitotic chromosomes were obtained from wing imaginal discs or testes of final instar larvae. Meiotic chromosomes in the pachytene stage of prophase I were obtained either from the testes of last instar larvae or from the ovaries of 3–5-day old pupae. Briefly, tissues were dissected in saline solution, swollen in hypotonic solution (75 mM KCl) for either 5 min (ovaries) or 15 min (testes and wing imaginal discs) and then fixed in Carnoy’s fixative (ethanol, chloroform, acetic acid, 6:3:1) for 10 min. Cells dissociated in 60% acetic acid were spread on a heating plate at 45°C. All chromosome preparations were passed through a graded ethanol series (70%, 80%, and 100%, 30 s each) and stored at –80°C.

Fluorescence in situ hybridization (FISH) with the (TTAGG)*_n_*telomeric probe and genomic in situ hybridization (GISH) were performed according to the procedure described by Yoshido et al. [74]. (TTAGG)*_n_* telomeric sequences were generated by non-template PCR following the protocol of Sahara et al. [75]. Male and female genomic DNAs (gDNAs) of *H. euphorbiae* were obtained separately from last instar larvae by standard phenol-chloroform extraction. DNA probes were labeled by nick translation using a mixture of DNase I and DNA polymerase I (both Thermo Fisher Scientific, Waltham, MA, USA) with either aminoallyl-dUTP-Cy3 or fluorescein-12-dUTP (both Jena Bioscience, Jena, Germany).

Chromosome preparations were removed from the freezer, passed through the graded ethanol series, air-dried and then denatured in 70% formamide in 2× SSC for 3.5 min at 70°C. For one preparation, the probe cocktail contained 500 ng of fluorescein-labeled female gDNA, 100 ng of Cy3-labeled telomeric probe, 3 μg of unlabeled sonicated male gDNA, and 25 μg of sonicated salmon sperm DNA (Sigma-Aldrich, St. Louis, MO, USA) in 10 μl hybridization buffer (50% formamide, 10% dextran sulfate in 2× SSC). Denaturation of the probe cocktail was performed for 5 min at 90°C. The preparations were examined under a Zeiss Axioplan 2 microscope (Carl Zeiss, Jena, Germany). Digital images were captured with an Olympus CCD monochrome camera XM10 equipped with cellSens 1.9 digital imaging software (Olympus Europa Holding, Hamburg, Germany) and processed with Adobe Photoshop CS4.

### Karyotype-based automated chromosome annotation and size estimation

The meiotic chromosome image was preprocessed using the image processing software GIMP (version 2.10) [76] to manually cut out individual chromosome bivalents. The processed image was loaded into the tool napari-karyotype (version c41103e) [49]. The image segmentation threshold, blur factor and genome size were set to 0.13, 0.5 and 504 Mb, respectively.

### Genome size estimation

Genome sizes of two hawkmoth species were estimated following the flow cytometry protocol with propidium iodide-stained nuclei as described previously [77]. Neural tissue from frozen (−80°C) adult samples of *H. vespertilio* and *H. euphorbiae* (from one head each; the same individuals used for the PacBio genome and Hi-C sequencing) and neural tissue from the internal reference standard *Acheta domesticus* (female, 1C = 2 Gb) were each chopped with a razor blade in a Petri dish containing 2 ml of ice-cold Galbraith buffer. The suspension was filtered through a 42-μm nylon mesh, then stained with the intercalating fluorochrome propidium iodide (PI, Thermo Fisher Scientific) and treated with Rnase A (Sigma-Aldrich), each at a final concentration of 25 μg/ml. The mean red PI fluorescence of the stained nuclei was quantified using a Beckman-Coulter CytoFLEX flow cytometer with a solid-state laser emitting at 488 nm. Fluorescence intensities of 10,000 nuclei per sample were recorded. Subsequently, the nuclei suspensions of *H. vespertilio* and *H. euphorbiae* were each mixed with the nuclei suspension of the internal reference standard (see above) and again the fluorescence intensities of 10,000 nuclei per mixed sample were recorded. For the histogram analyses, we used CytExpert 2.3 software. The total amount of DNA in each sample of the two *Hyles* species was calculated as the ratio of the mean fluorescence signal of the 2C peak of the stained nuclei of the respective species divided by the mean fluorescence signal of the 2C peak of the stained nuclei of the reference standard times the 1C amount of DNA in the reference standard. Three replicates, each from the same individual of *H. vespertilio* and *H. euphorbiae*, were measured on three different days to minimize possible random instrumental errors. Genome size was reported as 1C, the mean amount of DNA in Mb in a haploid nucleus.

### PacBio genome DNA and sequencing

Head tissue (38 mg) from a single male *Hyles euphorbiae* specimen (the same single individual used for genome size estimation) was used for high molecular weight DNA extraction using an adaptation of the protocol of Miller et al. [78]. Final purity and concentrations of DNA were measured using the NanoPhotometer (Implen GmbH, Munich, Germany) and the Qubit Fluorometer (Thermo Fisher Scientific, Waltham, MA). One SMRTbell library was constructed following the instructions of the SMRTbell Express Prep kit v2.0 with Low DNA Input Protocol (Pacific Biosciences, Menlo Park, CA). The total input DNA for the library was 3 µg. The library was loaded at an on-plate concentration of 80 pM using diffusion loading. One SMRT cell sequencing run was performed on the Sequel System II in CCS mode using 30-hour movie time with 2 hours of pre-extension and sequencing chemistry V2.0.

### Genome assembly of *Hyles euphorbiae*

We created PacBio CCS reads from the *Hyles euphorbiae* subreads.bam file using PacBio’s ccs command line tool (version 6.3.0, --all flag was used). We obtained 7.9 Gb of high-quality CCS reads (HiFi reads, rq > 0.99) with an N50 of 11.74 kb. To further increase the read coverage, we applied DeepConsensus (v0.2 with default settings, actc v0.1.1) [79] on all reads of the previous ccs step and obtained a total yield of 8.8 Gb (N50: 11.83 kb). We ran HiFiasm (version 0. 16.1-r375) [36] to create the contig assembly. Remaining haplotypic duplications in the primary contig set were removed using default parameters in the purge-dups pipeline (v.1.2.3) [80, 81]. The assembly was scaffolded with HiC data using yahs (v 1.1a) [82] and manually curated with higlass [83]. Remaining gaps in the scaffolds were filled by mapping the raw PacBio subreads with pbmm2 (version 1.7.0) [84], and for alignment piles that fully spanned the gap regions with 1000 anchor bases on both sides, a consensus sequence was produced with gcpp (version 2.0.2) [85]. The consensus sequence was used to fill a gap only if: 1) the complete consensus sequence was covered at least 5x; and 2) the coverage profile of the closed gaps fully supported the consensus sequence (i.e. no alignment breaks or huge repeat alignment piles occurred). Two rounds of error correction were performed by aligning the DeepConsensus reads to the assembly using pbmm2, calling variants using DeepVariant (version 1.3.0) [86] and correcting homozygous errors using bcftools [87, 88] consensus. The assembly was checked for contamination using blobtoolkit (version 1.1) [89] and an in-house pipeline that screens several blast databases. Assessment of completeness regarding single copy orthologs was performed via Benchmarking Using Single Copy Orthologues (BUSCO) (version 5.2.2) [90] together with the lepidoptera_odb10 set and the options “–long –offline”. and merqury (version 1.3) [91]. QV values were generated for the final scaffolds (*H. euphorbiae* QV=58.1). The mitochondrial genome was created using the mitoHifi pipeline (version 2) [45, 46] based on CCS reads and the closely related mitochondrial reference genome of *Theretra oldenlandiae* (NCBI accession: MN885801.1).

### Hi-C sequence data

Dovetail Hi-C libraries for *H. vespertilio* and *H. euphorbiae* were prepared from head tissues (52.2 mg and 40.6 mg; the same single individuals used for PacBio genome sequencing and genome size estimation) using the Dovetail Hi-C kit (Dovetail Genomics, Scotts Valley, CA, USA) following the manufacturer’s protocol (version 1.4) for insect samples. Briefly, chromatin was fixed with formaldehyde and then extracted. The fixed chromatin was digested with DpnII, the 5’ overhangs were filled with biotinylated nucleotides and the free blunt ends were ligated. After ligation, the crosslinks were reversed, the associated proteins were degraded and the DNA was purified. DNA was then sheared to a mean fragment size of ∼350 bp and sequencing libraries were generated using Illumina-compatible adapters. Biotinylated fragments were captured with streptavidin beads before PCR amplification.

The fragment size distribution and concentration of the final PacBio and Dovetail Hi-C libraries were assessed using the TapeStation (Agilent Technologies) and the Qubit Fluorometer (Thermo Fisher Scientific, Waltham, MA), respectively. The Hi-C libraries were sequenced on a NovaSeq 6000 platform at Novogene (UK), generating 100 million 2 × 150 bp paired-end reads each with a total volume of 30 Gb.

### Scaffolding the assembly of *H. vespertilio* with HiRise

The *H. vespertilio* input assembly of Pippel et al. [12] and the *H. vespertilio* Dovetail Hi-C library reads were used as input data for HiRise, a software pipeline specifically designed for using proximity ligation data to scaffold genome assemblies [40]. The Dovetail Hi-C library sequences were aligned to the draft input assembly using a modified SNAP read mapper (http://snap.cs.berkeley.edu). The separations of the Hi-C read pairs mapped within draft scaffolds were analyzed using HiRise version 2.1.7 to produce a likelihood model for the genomic distance between the read pairs. The model was used to identify and break putative misjoins, score prospective joins, and make joins above a threshold.

### Annotation and methylation profile

Structural annotation of protein coding genes was performed using TOGA [92], a method that uses pairwise genome alignment chains between an annotated reference genome (here *Manduca sexta*) and other query species (here *Hyles euphorbiae* and *H. vespertilio*). Briefly, TOGA uses machine learning to infer orthologous loci for each reference transcript, using the concept that orthologous genes display more alignments between intronic and flanking intergenic regions. TOGA then projects each reference transcript to its orthologous query locus using CESAR 2.0 [93], a hidden Markov model method that takes reading frame and splice site annotation of the reference exons into account. CESAR avoids spurious frameshifts and is able to detect evolutionary splice site shifts and precise intron deletions [93, 94]. Using the CESAR alignment, TOGA determines whether the transcript has inactivating mutations (frameshifting mutations, premature stop codons, splice site disrupting mutations, deletions of entire coding exons).

To generate whole genome alignments as input for TOGA, we aligned the assemblies of *H. euphorbiae* and *H. vespertilio* to the *Manduca sexta* (GCF_014839805.1) [37] assembly. To compare contiguity between *H. euphorbiae*, *H. vespertilio*, *L. populi*, *Hemaris fuciformis*, *D. porcellus*, *Mimas tiliae* and *Manduca Sexta*, Quast 5.0.2 [95] was used.

Genomes were aligned using LASTZ (version 1.04.03) [96] with the parameters (K = 2400, L = 3000, Y = 9400, H = 2000 and the lastz default scoring matrix). Then, we used axtChain [47] (default parameters except linearGap=loose) to compute co-linear alignment chains, RepeatFiller [97] (default parameters) to capture previously missed alignments between repetitive regions and chainCleaner [98] (default parameters except minBrokenChainScore=75000 and -doPairs) to improve alignment specificity. *H. vespertilio* was used as reference and *H. euphorbiae*, *B. mori* and *M. sexta* as queries. We used the NCBI annotation (GCF_014839805.1_JHU_Msex_v1.0_genomic.gff.gz) as input for TOGA. The mitochondrial genome of *H. euphorbiae* was annotated using the MITOS WebServer [99] and the result illustrated using shinyCircos [100]. This software was also used for the CIRCOS-Plots of the aligned genomes.

### Genome alignments

With the aim of naming the chromosomes according to homology to *B. mori* Nucleotide homology was assessed based on sequence alignment of *H. vespertilio*, *H. euphorbiae* and *M. sexta* contigs via scaffold chaining as the primary tool for discovering similarities using LASTZ [96]. Visualization was carried out using CIRCOS plots (Fig. 4) using positional information of synteny blocks (in the bed files) of the genome alignments as input (c.f. Table S3).

The *Hyles euphorbiae* and *H. vespertilio* genome assemblies (with annotations and probabilities of site modification/methylation, and genome alignments, see below) are accessible in the Senckenberg Genome Browser (https://genome.senckenberg.de/cgi-bin/hgTracks?db=HLhylEup1; https://genome.senckenberg.de/cgi-bin/hgTracks?db=HLhylVes2). A genome alignment of *H. vespertilio* to *B. mori* (GCF_014905235.1 [41]) was generated as above and used to postulate chromosome homologies and name chromosomes accordingly. Detailed values of the proportions of homologous regions per chromosome are provided in Table S1.

The genomes of *Mimas tiliae* [14], *Laothoe populi* [15], *Hemaris fuciformis* [38] and *Deilephila porcellus* [39] (these are the additional genomes of Sphingidae that were available on 14.10.2022) were included for assembly statistics comparisons and a phylogenetic comparison of repeat evolution.

### Analyses of repeat and transposon content

Comparative annotation of repeats for *H. vespertilio*, *H. euphorbiae*, *D. porcellus, Mimas tiliae, L. populi, H. fuciformis,* and *Manduca sexta* was performed using RepeatModeler (version 2.0.3) [101] and RepeatMasker (version 4.1.2-p1) [102] as provided in Dfam TETools docker container (version 1.85) [103], in combination with rmblastn (version 2.11.0+). The repeat library for RepeatMasker contains a combination of all repeat families identified with RepeatModeler, combined and made non-redundant using Cdhit (version 4.8.1) [104] and manual curation of models according to guidelines detailed in [105]. Occurrence of various transposable element families were illustrated on the respective branches of a tree drawn by hand based on Kawahara and Barber [44], in relative order according to the assembly divergence and sequence amount as provided by a helper script of repeatmasker (calcDivergenceFromAlign.pl). TE families with higher sequence diversity (presumed to represent older transposon bursts) are drawn more left on the branches, TE families shared between taxa (and with high sequence diversity) are drawn on the branch of their common ancestors.

### Methylation profile

For the methylation profile, the *H. euphorbiae* PacBio HiFi reads containing 5mC base modification tags were converted to CpG/5mC data using pb-CpG-tools (version 1.1) [106] and uploaded to the genome browser (s.a.). Sites with a modification probability of <50% are marked blue and >50% red.

### Chromosome structure analysis

To compare chromosome structure across species, single copy orthologs were inferred using BUCSOs (version 5.4) (‘lepidoptera odb 10’ dataset, metaeuk mode) [107]. Orthologues were filtered to retain those that can be assigned to a Merian element, using the table of orthologues assignments to Merian elements from Wright *et al*. [42]. Next, unlocalized scaffolds were removed by only retaining scaffolds with three or more single copy orthologs. Finally, synteny between pairs of genomes was then visualized using Oxford plots with using custom code [108]. which plots the relative position of each ortholog along the chromosomes of each genome. In the resulting Oxford plots, orthologues were colored by Merian element identity to identify chromosomes resulting from fusion events. Chromosomes resulting from fusion events were also verified by running https://github.com/charlottewright/lep_fusion_fission_finder with a window size of 17. This identifies chromosomes which are the product of fusion events based on the presence of orthologues that correspond to more than one Merian element within a single chromosome.

### Wing pattern genes

The positions of the wing pattern genes *optix, WntA*, *cortex, aristaless* and *distal-less* (see Table S2 for accession numbers of references) were identified in the two *Hyles* genomes using the online BLAT tool [47, 48] with default options as implemented in the Senckenberg Genome browser. The BLAT results are presented sorted by alignment length, with the longest selected for each gene.

For the *cortex* gene, we additionally compared the *Hyles* data with the sequences of *Biston betularia* (Geometridae; KT182637), in which the common pale (*typica*) form was replaced by a previously unknown black (*carbonaria*) form during the Industrial Revolution, driven by the interaction between bird predation and smoke pollution 1. [109] and caused by a transposon insertion in a *cortex* intron [32, 110].

## Declarations

### Ethics approval and consent to participate

The individual of the protected species *H. euphorbiae* was collected with an exception issued in a permit by “Landratsamt Meißen, Kreisumweltamt, Untere Naturschutzbehörde” (AZ 364.621-377/2009-4026/2017-45577/2021) for the purpose of scientific study.

#### Consent for publication

not applicable

### Availability of data and materials

The data generated and analyzed during the current study are available in the NCBI repository, [https://www.ncbi.nlm.nih.gov/genome/?term=hyles]. The SRA accession number for *H. euphorbiae* PacBio Sequel HiFi is SRX13604162, and for the Dovetail Hi-C data is SRX14310646. The assembled genome is accessible under GCA_023078785.2, and accession numbers are itemized in the following. Mitochondrion: CM041059, chromosome Z: CM041058 and chromosomes 2-29: CM041030-CM041057).

The SRA accession number for the Dovetail *Hyles vespertilio* Hi-C raw data is SRX14530528. The new assembly presented in this work is accessible under GCA_009982885.2, and accession numbers are itemized in the following. Chromosome Z: CM042833 and chromosomes 2-29: CM042805-CM042832.

### Authors’ contributions

AKH conceived the study and partnerships, did field work, performed analyses, produced and edited figures, and wrote the text. TS supported assembly, scaffolding, annotation, wrote, and edited text. FP, CW & LP proposed and performed analyses, produced figures, and wrote text. AY & FM contributed karyotype analyses with figures and text (with additional editing of the entire ms). HD contributed field and laboratory work and SW & CG took responsibility for DNA isolation, sequencing, and contributed text. MH provided bioinformatics infrastructure and supervision, scripts, intermediate results, advice, and performed alignments, MP performed assembly, scaffolding, numerous analyses and provided the basis for figures. All authors read and approved the final manuscript.

## Acknowledgements

This study benefitted from the sharing of expertise within the DFG priority program SPP 1991 Taxon-Omics. Dovetail Genomics provided HiRise scaffolding service. Thanks go to Alexander Ben Hamadou (LOEWE-TBG) for DNA isolation and library preparations. We thank Ian J. Kitching for language correction and helpful suggestions.

## References

1. Hundsdoerfer AK, Päckert M, Kehlmaier C, Strutzenberger P, Kitching IJ: Museum archives revisited: Central Asiatic hawkmoths reveal exceptionally high late Pliocene species diversification (Lepidoptera, Sphingidae). Zoologica Scripta 2017, 46(5):552–570.

2. Hundsdoerfer AK, Kitching IJ, Wink M: A molecular phylogeny of the hawkmoth genus *Hyles* (Lepidoptera: Sphingidae, Macroglossinae). Molecular Phylogenetics and Evolution 2005, 35:442–458.

3. Hundsdoerfer AK, Lee KM, Kitching IJ, Mutanen M: Genome-wide SNP data reveal an overestimation of species diversity in a group of hawkmoths. Genome Biology and Evolution 2019, 11(8):2136–2150.

4. Hundsdoerfer AK, Tshibangu JN, Wetterauer B, Wink M: Sequestration of phorbol esters by aposematic larvae of *Hyles euphorbiae* (Lepidoptera: Sphingidae)? Chemoecology 2005, 15(4):261–267.

5. Hundsdoerfer AK, Rubinoff D, Attié M, Kitching IJ, Wink M: A revised molecular phylogeny of the globally distributed hawkmoth genus *Hyles* (Lepidoptera: Sphingidae), based on mitochondrial and nuclear DNA sequences. Molecular Phylogenetics and Evolution 2009, 52:852–865.

6. Hundsdoerfer AK, Buchwalder K, O’Neill MA, Dobler S: Chemical ecology traits in an adaptive radiation: TPA-sensitivity and detoxification in *Hyles* and *Hippotion* (Sphingidae, Lepidoptera) larvae. Chemoecology 2019, 29(1):35–47.

7. Mende MB, Bartel M, Hundsdoerfer AK: A comprehensive phylogeography of the *Hyles euphorbiae* complex (Lepidoptera: Sphingidae) indicates a ’glacial refuge belt’. Scientific Reports 2016, 6:29527

8. Les Sphingidae de France [http://sphingidae-haxaire.com/index.php/macroglossinae-2/hyles-vespertilio/]

9. Hundsdoerfer AK, Kitching IJ: Morphological evolution in *Hyles* Hübner, 1819 hawkmoths (Lepidoptera, Sphingidae): reconstructing the ancestral *Hyles* habitus. Nota Lepidopterologica 2020, 43:181.

10. Danner F, Eitschberger U, Surholt B: Die Schwärmer der westlichen Palaearktis. Bausteine zu einer Revision (Lepidoptera: Sphingidae). Herbipoliana 1998, 4:1–368 (Textband), 361–720 (Tafelband).

11. Hundsdoerfer AK, Kitching IJ: Ancient incomplete lineage sorting of Hyles and Rhodafra (Lepidoptera: Sphingidae). Organisms Diversity & Evolution 2020, 20(3):527–536.

12. Pippel M, Jebb D, Patzold F, Winkler S, Vogel H, Myers G, Hiller M, Hundsdoerfer AK: A highly contiguous genome assembly of the bat hawkmoth Hyles vespertilio (Lepidoptera: Sphingidae). GigaScience 2020, 9(1):giaa001.

13. Patzold F, Zilli A, Hundsdoerfer AK: Advantages of an easy-to-use DNA extraction method for minimal-destructive analysis of collection specimens. PLoS One 2020, 15(7):e0235222.

14. Boyes D, Holland P, University of Oxford and Wytham Woods Genome Acquisition Lab L, al. e: The genome sequence of the lime hawk-moth, Mimas tiliae (Linnaeus, 1758). Wellcome Open Research 2021, 6.

15. Boyes D, Holland PW, Consortium DToL: The genome sequence of the poplar hawk-moth, Laothoe populi (Linnaeus, 1758). Wellcome Open Research 2021 [version 1 peer review: 1 approved, 1 approved with reservations], 6:237.

16. Beldade P, Brakefield PM: The genetics and evo-devo of butterfly wing patterns. Nat Rev Genet 2002, 3(6):442–452.

17. Nijhout HF: Elements of butterfly wing patterns. J Exp Zool 2001, 291(3):213–225.

18. Beldade P, Monteiro A: Eco-evo-devo advances with butterfly eyespots. Curr Opin Genet Dev 2021, 69:6–13.

19. Monteiro A, Glaser G, Stockslager S, Glansdorp N, Ramos D: Comparative insights into questions of lepidopteran wing pattern homology. Bmc Dev Biol 2006, 6.

20. Van Belleghem SM, Rastas P, Papanicolaou A, Martin SH, Arias CF, Supple MA, Hanly JJ, Mallet J, Lewis JJ, Hines HM et al: Complex modular architecture around a simple toolkit of wing pattern genes. Nat Ecol Evol 2017, 1(3):52.

21. Dhungel B, Ohno Y, Matayoshi R, Iwasaki M, Taira W, Adhikari K, Gurung R, Otaki JM: *distal-less* induces elemental color patterns in *Junonia* butterfly wings. Zool Lett 2016, 2.

22. Connahs H, Rhen T, Simmons RB: Transcriptome analysis of the painted lady butterfly, *Vanessa cardui* during wing color pattern development. Bmc Genomics 2016, 17.

23. Martin A, Reed RD: *wingless* and *aristaless2* define a developmental ground plan for moth and butterfly wing pattern evolution. Molecular Biology and Evolution 2010, 27(12):2864–2878.

24. Stevens M, Merilaita S: Animal camouflage: current issues and new perspectives. Philosophical Transactions of the Royal Society B: Biological Sciences 2009, 364(1516):423–427.

25. Hiyama A, Taira W, Otaki JM: Color-pattern evolution in response to environmental stress in butterflies. Frontiers in Genetics 2012, 3(15).

26. Bastide H, Saenko SV, Chouteau M, Joron M, Llaurens V: Dominance mechanisms in supergene alleles controlling butterfly wing pattern variation: insights from gene expression in *Heliconius numata*. Heredity 2023, 130(2):92–98.

27. Jay P, Leroy M, Le Poul Y, Whibley A, Arias M, Chouteau M, Joron M: Association mapping of colour variation in a butterfly provides evidence that a supergene locks together a cluster of adaptive loci. Philos T R Soc B 2022, 377(1856).

28. Zhang L, Mazo-Vargas A, Reed RD: Single master regulatory gene coordinates the evolution and development of butterfly color and iridescence. Proc Natl Acad Sci U S A 2017, 114(40):10707–10712.

29. Livraghi L, Martin A, Gibbs M, Braak N, Arif S, Breuker CJ: CRISPR/Cas9 as the key to unlocking the secrets of butterfly wing pattern development and its evolution. In: Advances in Insect Physiology. vol. 54: Elsevier; 2018: 85–115.

30. Jiggins CD, Wallbank RW, Hanly JJ: Waiting in the wings: what can we learn about gene co-option from the diversification of butterfly wing patterns? Philos Trans R Soc Lond B Biol Sci 2017, 372(1713).

31. Nadeau NJ, Pardo-Diaz C, Whibley A, Supple MA, Saenko SV, Wallbank RW, Wu GC, Maroja L, Ferguson L, Hanly JJ et al: The gene cortex controls mimicry and crypsis in butterflies and moths. Nature 2016, 534(7605):106–110.

32. van’t Hof AE, Campagne P, Rigden DJ, Yung CJ, Lingley J, Quail MA, Hall N, Darby AC, Saccheri IJ: The industrial melanism mutation in British peppered moths is a transposable element. Nature 2016, 534(7605):102–105.

33. Brunetti CR, Selegue JE, Monteiro A, French V, Brakefield PM, Carroll SB: The generation and diversification of butterfly eyespot color patterns. Curr Biol 2001, 11(20):1578–1585.

34. Westerman EL, VanKuren NW, Massardo D, Tenger-Trolander A, Zhang W, Hill RI, Perry M, Bayala E, Barr K, Chamberlain N et al: aristaless controls butterfly wing color variation used in mimicry and mate choice. Curr Biol 2018, 28(21):3469.

35. Bayala EX, VanKuren N, Massardo D, Kronforst MR: *aristaless1* has a dual role in appendage formation and wing color specification during butterfly development. BMC Biology 2023, 21:1–19.

36. Cheng H, Concepcion GT, Feng X, Zhang H, Li H: Haplotype-resolved de novo assembly using phased assembly graphs with hifiasm. Nature methods 2021, 18(2):170–175.

37. Gershman A, Romer TG, Fan Y, Razaghi R, Smith WA, Timp W: De novo genome assembly of the tobacco hornworm moth (Manduca sexta). G3 2021, 11(1):1–9.

38. Mulhair PO, Holland PW: Evolution of the insect Hox gene cluster: Comparative analysis across 243 species. Seminars in Cell & Developmental Biology 2022, https://doi.org/10.1016/j.semcdb.2022.11.010.

39. Boyes D, University of Oxford and Wytham Woods Genome Acquisition Lab L, Darwin Tree of Life Barcoding collective c, al. e: The genome sequence of the small elephant hawk moth, Deilephila porcellus (Linnaeus, 1758) Wellcome Open Research 2022, 7:80.

40. Putnam NH, O’Connell BL, Stites JC, Rice BJ, Blanchette M, Calef R, Troll CJ, Fields A, Hartley PD, Sugnet CW: Chromosome-scale shotgun assembly using an in vitro method for long-range linkage. Genome Research 2016, 26(3):342–350.

41. Kawamoto M, Jouraku A, Toyoda A, Yokoi K, Minakuchi Y, Katsuma S, Fujiyama A, Kiuchi T, Yamamoto K, Shimada T: High-quality genome assembly of the silkworm, Bombyx mori. Insect Biochemistry and Molecular Biology 2019, 107:53–62.

42. Wright CJ, Stevens L, Mackintosh A, Lawniczack M, Blaxter M: Chromosome evolution in Lepidoptera. BioRxiv 2023, https://www.biorxiv.org/content/10.1101/2023.05.12.540473v1.

43. Yasukochi Y, Tanaka-Okuyama M, Shibata F, Yoshido A, Marec F, Wu C, Zhang H, Goldsmith MR, Sahara K: Extensive conserved synteny of genes between the karyotypes of Manduca sexta and Bombyx mori revealed by BAC-FISH mapping. PLoS One 2009, 4(10):e7465.

44. Kawahara AY, Barber JR: Tempo and mode of antibat ultrasound production and sonar jamming in the diverse hawkmoth radiation. Proceedings of the National Academy of Sciences 2015, 112(20):6407–6412.

45. Allio R, Schomaker-Bastos A, Romiguier J, Prosdocimi F, Nabholz B, Delsuc F: MitoFinder: Efficient automated large-scale extraction of mitogenomic data in target enrichment phylogenomics. https://github.com/marcelauliano/MitoHiFi. Mol Ecol Resour 2020, 20(4):892–905.

46. Uliano-Silva M, Gabriel R, Ferreira J, Krasheninnikova K, Consortium DToL, Formenti G, Abueg L, …, McCarthy SA: MitoHiFi: a python pipeline for mitochondrial genome assembly from PacBio High Fidelity reads. bioRxiv 2022, 2022–12.

47. Kent WJ, Baertsch R, Hinrichs A, Miller W, Haussler D: Evolution’s cauldron: duplication, deletion, and rearrangement in the mouse and human genomes. Proceedings of the National Academy of Sciences of the United States of America 2003, 100(20):11484–11489.

48. Lee BT, Barber GP, Benet-Pages A, Casper J, Clawson H, Diekhans M, Fischer C, Gonzalez JN, Hinrichs AS, Lee CM et al: The UCSC Genome Browser database: 2022 update. Nucleic Acids Res 2022, 50(D1):D1115–D1122.

49. napari, contributors: napari: a multi-dimensional image viewer for python. In.; 2019.

50. Hundsdoerfer AK, Kitching IJ, Wink M: The phylogeny of the *Hyles euphorbiae*-complex (Lepidoptera: Sphingidae): molecular evidence from sequence data and ISSR-PCR fingerprints. Organisms Diversity and Evolution 2005, 5:173–198.

51. Hanrahan SJ, Johnston JS: New genome size estimates of 134 species of arthropods. Chromosome Research 2011, 19(6):809–823.

52. d’Alencon E, Sezutsu H, Legeai F, Permal E, Bernard-Samain S, Gimenez S, Gagneur C, Cousserans F, Shimomura M, Brun-Barale A et al: Extensive synteny conservation of holocentric chromosomes in Lepidoptera despite high rates of local genome rearrangements. Proc Natl Acad Sci U S A 2010, 107(17):7680–7685.

53. Lavoie CA, Platt RN, Novick PA, Counterman BA, Ray DA: Transposable element evolution in *Heliconius* suggests genome diversity within Lepidoptera. Mob DNA 2013, 4(1):21.

54. Lynch M, Walsh B: The origins of genome architecture vol. 98. Sunderland, MA: Sinauer Associates; 2007.

55. Shah A, Hoffman JI, Schielzeth H: Comparative analysis of genomic repeat content in Gomphocerine grasshoppers reveals expansion of satellite DNA and helitrons in species with unusually large genomes. Genome Biology and Evolution 2020, 12(7):1180–1193.

56. Hollister JD, Gaut BS: Epigenetic silencing of transposable elements: a trade-off between reduced transposition and deleterious effects on neighboring gene expression. Genome Research 2009, 19(8):1419–1428.

57. Charlesworth B, Sniegowski P, Stephan W: The evolutionary dynamics of repetitive DNA in eukaryotes. Nature 1994, 371(6494):215–220.

58. Talla V, Suh A, Kalsoom F, Dinca V, Vila R, Friberg M, Wiklund C, Backstrom N: Rapid increase in genome size as a consequence of transposable element hyperactivity in Wood-White (Leptidea) Butterflies. Genome Biology and Evolution 2017, 9(10):2491–2505.

59. Ellegren H, Smeds L, Burri R, Olason PI, Backstrom N, Kawakami T, Kunstner A, Makinen H, Nadachowska-Brzyska K, Qvarnstrom A et al: The genomic landscape of species divergence in Ficedula flycatchers. Nature 2012, 491(7426):756–760.

60. Feliciello I, Akrap I, Brajkovic J, Zlatar I, Ugarkovic D: Satellite DNA as a driver of population divergence in the red flour beetle Tribolium castaneum. Genome Biol Evol 2014, 7(1):228–239.

61. Maumus F, Fiston-Lavier AS, Quesneville H: Impact of transposable elements on insect genomes and biology. Curr Opin Insect Sci 2015, 7:30–36.

62. Pittaway AR: The hawkmoths of the Western Palaearctic. Colchester: Harley Books; 1993.

63. Brower AVZ: Rapid morphological radiation and convergence among races of the butterfly *Heliconius erato* inferred from patterns of mitochondrial DNA evolution. Proceedings of the National Academy of Science of the United States of America 1994, 91:6491–6495.

64. Pringle EG, Baxter SW, Webster CL, Papanicolaou A, Lee SF, Jiggins CD: Synteny and chromosome evolution in the lepidoptera: evidence from mapping in Heliconius melpomene. Genetics 2007, 177(1):417–426.

65. Yasukochi Y, Ashakumary LA, Baba K, Yoshido A, Sahara K: A second-generation integrated map of the silkworm reveals synteny and conserved gene order between lepidopteran insects. Genetics 2006, 173(3):1319–1328.

66. Ahola V LR, Somervuo P, Salmela L, Koskinen P, Rastas P et al.: The Glanville fritillary genome retains an ancient karyotype and reveals selective chromosomal fusions in Lepidoptera. Nature Communications 2014(5):4737.

67. Sahara K, Yoshido A, Shibata F, Fujikawa-Kojima N, Okabe T, Tanaka-Okuyama M, Yasukochi Y: FISH identification of Helicoverpa armigera and Mamestra brassicae chromosomes by BAC and fosmid probes. Insect Biochem Mol Biol 2013, 43(8):644–653.

68. van’t Hof AE, Nguyen P, Dalikova M, Edmonds N, Marec F, Saccheri IJ: Linkage map of the peppered moth, *Biston betularia* (Lepidoptera, Geometridae): a model of industrial melanism. Heredity 2013, 110(3):283–295.

69. Yoshido A, Bando H, Yasukochi Y, Sahara K: The Bombyx mori karyotype and the assignment of linkage group. Genetics 2005, 170:675–685.

70. Jiggins CD: What can we learn about adaptation from the wing pattern genetics of Heliconius butterflies? In: Diversity and evolution of butterfly wing patterns. Springer, Singapore; 2017: 173–188.

71. Reed RD, Papa R, Martin A, Hines HM, Counterman BA, Pardo-Diaz C, Jiggins CD, Chamberlain NL, Kronforst MR, Chen R et al: optix drives the repeated convergent evolution of butterfly wing pattern mimicry. Science 2011, 333(6046):1137–1141.

72. Maddison WP: Gene trees in species trees. Systematic Biology 1997, 46(3):523–536.

73. Yoshido A, Sahara K, Yasukochi Y, Sharakhov I: Silk moths (Lepidoptera). In: Protocols for Cytogenetic Mapping of Arthropod Genomes Edited by Sharakhov IV. Boca Ranton, FL, USA: CRC Press; 2015: 219–256.

74. Yoshido A, Marec F, Sahara K: Resolution of sex chromosome constitution by genomic in situ hybridization and fluorescence in situ hybridization with (TTAGG)(n) telomeric probe in some species of Lepidoptera. Chromosoma 2005, 114(3):193–202.

75. Sahara K, Marec F, Traut W: TTAGG telomeric repeats in chromosomes of some insects and other arthropods. Chromosome Research 1999, 7(6):449–460.

76. Team TGD: GIMP. Retrieved from https://www.gimp.org. In.; 2019.

77. Hare EE, Johnston JS: Genome size determination using flow cytometry of propidium iodide-stained nuclei. In: Molecular methods for evolutionary genetics. Springer; 2012: 3–12.

78. Miller S, Dykes D, Polesky H: A simple salting out procedure for extracting DNA from human nucleated cells. Nucleic Acids Research 1988, 16(3):1215.

79. Align subreads to ccs reads [https://github.com/PacificBiosciences/actc]

80. Guan D, McCarthy SA, Wood J, Howe K, Wang Y, Durbin R: Identifying and removing haplotypic duplication in primary genome assemblies. Bioinformatics 2020, 36(9):2896–2898.

81. Haplotypic duplication identification tool [https://github.com/dfguan/purge_dups#step-1-run-minimap2-to-align-pacbio-data-and-generate-paf-files-then-calculate-read-depth-histogram-and-base-level-read-depth-commands-are-as-follows]

82. Zhou C, McCarthy SA, Durbin R: YaHS: yet another Hi-C scaffolding tool. bioRxiv 2022.

83. Kerpedjiev P, Abdennur N, Lekschas Fea: HiGlass: web-based visual exploration and analysis of genome interaction maps. Genome Biology 2018, 19:125.

84. A minimap2 frontend for PacBio native data formats [https://github.com/PacificBiosciences/pbmm2]

85. GCpp Generate Highly Accurate Reference Contigs

86. Poplin R, Chang P-C, Alexander D, Schwartz S, Colthurst T, Ku A, Newburger D, Dijamco J, Nguyen N, Afshar PT: A universal SNP and small-indel variant caller using deep neural networks. Nature Biotechnology 2018, 36(10):983–987.

87. bcftools - utilities for variant calling and manipulating VCFs and BCFs [https://samtools.github.io/bcftools/bcftools.html]

88. Li H, Handsaker B, Wysoker A, Fennell T, Ruan J, Homer N, Marth G, Abecasis G, Durbin R: The sequence alignment/map format and SAMtools. Bioinformatics 2009, 25(16):2078–2079.

89. Challis R, Richards E, Rajan J, Cochrane G, Blaxter M: BlobToolKit–interactive quality assessment of genome assemblies. G3: Genes, Genomes, Genetics 2020, 10:1361–1374.

90. Manni M, Berkeley MR, Seppey M, Simao FA, Zdobnov EM: BUSCO update: novel and streamlined workflows along with broader and deeper phylogenetic coverage for scoring of eukaryotic, prokaryotic, and viral genomes. arXiv preprint arXiv:210611799 2021.

91. Rhie A, Walenz BP, Koren S, Phillippy AM: Merqury: reference-free quality, completeness, and phasing assessment for genome assemblies. Genome Biology 2020, 21(1):1–27.

92. Kirilenko B, Munegowda C, Osipova E, Jebb D, Sharma V, Blumer M, Morales A, Ahmed A, Kontopoulos D, Hilgers L et al: TOGA integrates gene annotation with orthology inference at scale. under review.

93. Sharma V, Schwede P, Hiller M: CESAR 2.0 substantially improves speed and accuracy of comparative gene annotation. Bioinformatics 2017, 33(24):3985–3987.

94. Sharma V, Elghafari A, Hiller M: Coding exon-structure aware realigner (CESAR) utilizes genome alignments for accurate comparative gene annotation. Nucleic Acids Res 2016, 44(11):e103.

95. Gurevich A, Saveliev V, Vyahhi N, Tesler G: QUAST: quality assessment tool for genome assemblies. Bioinformatics 2013, 29(8):1072–1075.

96. Harris RS: Improved pairwise alignment of genomic DNA. A thesis in computer science and engineering.: The Pennsylvania State University; 2007.

97. Osipova E, Hecker N, Hiller M: RepeatFiller newly identifies megabases of aligning repetitive sequences and improves annotations of conserved non-exonic elements. bioRxiv 2019:696922.

98. Suarez HG, Langer BE, Ladde P, Hiller M: chainCleaner improves genome alignment specificity and sensitivity. Bioinformatics 2017, 33(11):1596–1603.

99. Bernt M, Donath A, Juhling F, Externbrink F, Florentz C, Fritzsch G, Putz J, Middendorf M, Stadler PF: MITOS: improved de novo metazoan mitochondrial genome annotation. Mol Phylogenet Evol 2013, 69(2):313–319.

100. Yu Y, Ouyang Y, Yao W: shinyCircos: an R/Shiny application for interactive creation of Circos plot. Bioinformatics 2018, 34(7):1229–1231.

101. Smit A, Hubley R, Green P: RepeatModeler Open-1.0. 2008–2015. Institute for Systems Biology, Seattle, USA Available from: http://www.repeatmaskerorg 2015.

102. Smit A, Hubley R, Green P: RepeatMasker Open-4.0. 2013–2015. Available from: http://www.repeatmasker.org. In.; 2015.

103. TETools [https://github.com/Dfam-consortium/TETools]

104. Fu L, Niu B, Zhu Z, Wu S, Li W: CD-HIT: accelerated for clustering the next-generation sequencing data. Bioinformatics 2012, 28:3150–3152.

105. Goubert C, Craig RJ, Bilat AF, Peona V, Vogan AA, Protasio AV: A beginner’s guide to manual curation of transposable elements. Mobile DNA. Mobile DNA 2022, 13:1–19.

106. pb-CpG-tools [https://github.com/PacificBiosciences/pb-CpG-tools]

107. Simao Neto F, Waterhouse R, Ioannidis P, Kriventseva E, Zdobnov E: BUSCO: assessing genome assembly and annotation completeness with single-copy orthologs. Bioinformatics 2015, 31:3210–3212.

108. Scripts associated with Hundsdoerfer, et al, 2023 [https://github.com/charlottewright/hyles_manuscript]

109. Cook LM: The rise and fall of the Carbonaria form of the peppered moth. Q Rev Biol 2003, 78(4):399–417.

110. van’t Hof AE, Edmonds N, Dalíková M, Marec F, Saccheri IJ: Industrial melanism in British peppered moths has a singular and recent mutational origin. Science 2011, 332(6032):958–960.

