## Supplementary Table S2 for "High-quality haploid genomes corroborate 29 chromosomes and highly conserved synteny of genes in *Hyles* hawkmoths (Lepidoptera: Sphingidae)"

**Accession numbers of reference wing pattern sequences and data type used for blat in the genome browser.** *Manduca sexta* *WntA* and *optix* are not displayed on the NCBI web portal.

| **Gene** | **Reference species** | **Accession number** | **BLAT** |
| --- | --- | --- | --- |
| *WntA* | *Heliconius himera* | AFC75686.1 | Protein sequence |
| *optix* | *Heliconius erato* | KC469894.1 | Protein sequence |
| *cortex* (isoform X1) | *Manduca sexta* | XM_030179809 | mRNA sequence |
| *aristaless* | *Manduca sexta* | XM_030185306.2, XM_037442080.1- XM_037442082.1 | mRNA sequences |
| *distal-less* | *Manduca sexta* | AY616435.1 | mRNA sequence |
