## Supplementary Figures for "High-quality haploid genomes corroborate 29 chromosomes and highly conserved synteny of genes in *Hyles* hawkmoths (Lepidoptera: Sphingidae)"

**Hundsdoerfer et al *Hyles* chromosome level: Supplementary figures**

**Fig. S1**

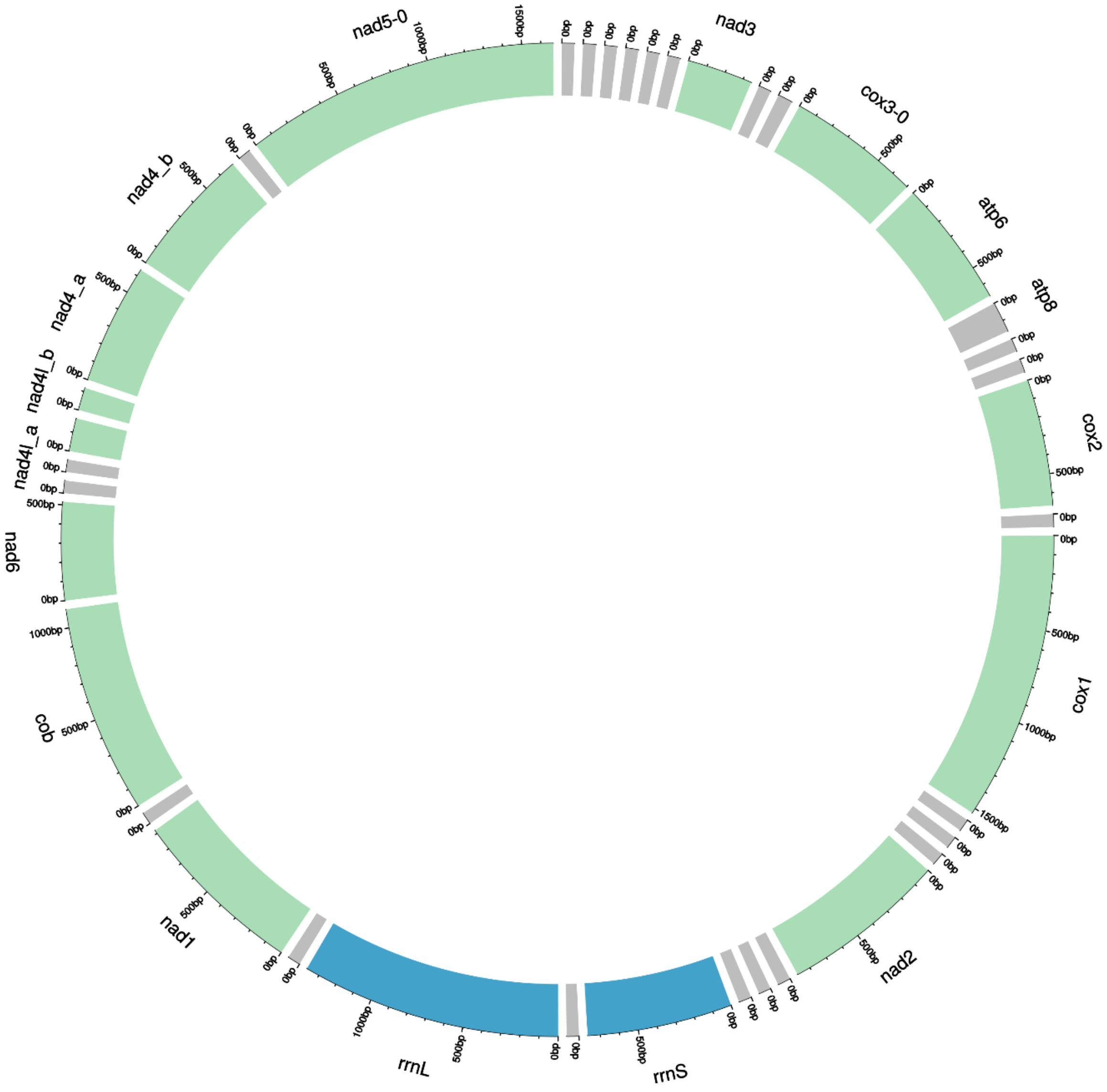


**The annotated mitochondrial genome of *Hyles euphorbiae*.**

It contains 13 protein coding genes, two ribosomal RNA (rRNA) genes plus 22 transfer RNA (tRNA) sequences and the control region.

**Fig. S2**

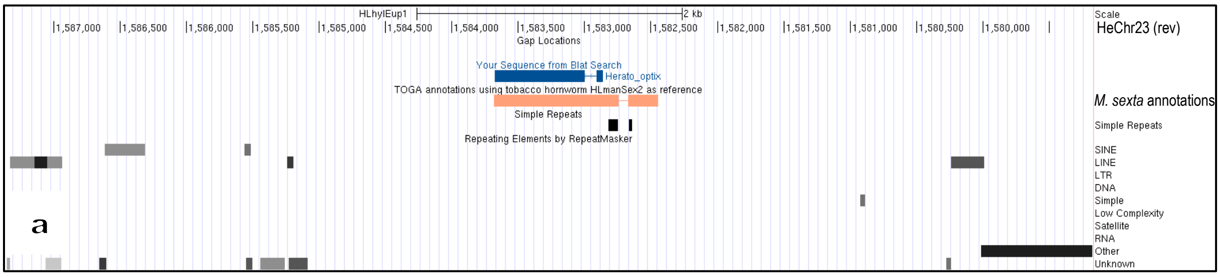

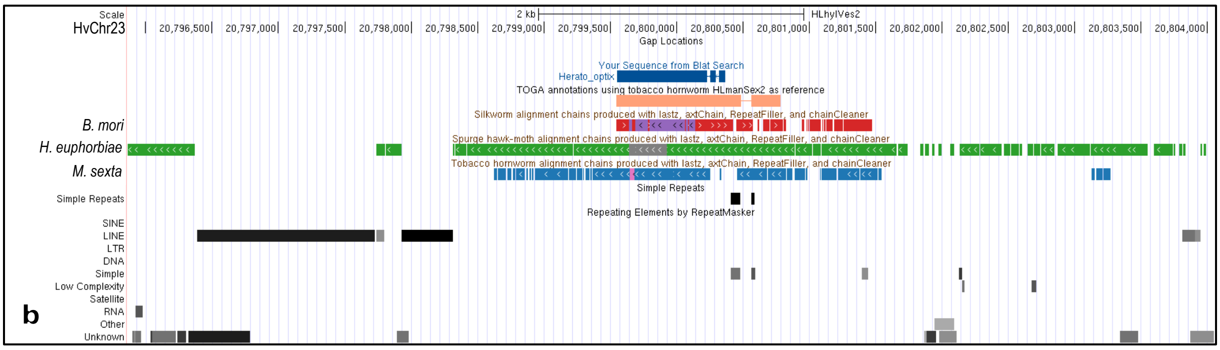

**BLAT result of the *Heliconius optix* protein sequence (KC469894) from the genome browser on chromosome 23 showing the *Hyles* genome alignment, annotation, and repeat occurrence in the vicinity in a) *H. euphorbiae* (inverted) and b) *H. vespertilio*.**

The *Heliconius erato optix* amino acid sequence BLAT search resulted in a match on chromosome 23 in both *Hyles* species (*B. mori* and *M. sexta* are included in b) for comparison).

The annotation track (bright orange) shows positions of exons annotated with TOGA, based on *M. sexta*.

Other sequence colors in b) are red and purple (main scaffold of *B. mori*), green (main scaffold of *H. euphorbiae,* inverted), blue (main scaffold of *M. sexta*). Additional scaffolds are given yet another color to be able to distinguish which chains align which scaffold. The darker the shades of grey of the repeats, the closer they are to the repeat consensus sequence.

**Fig. S3**

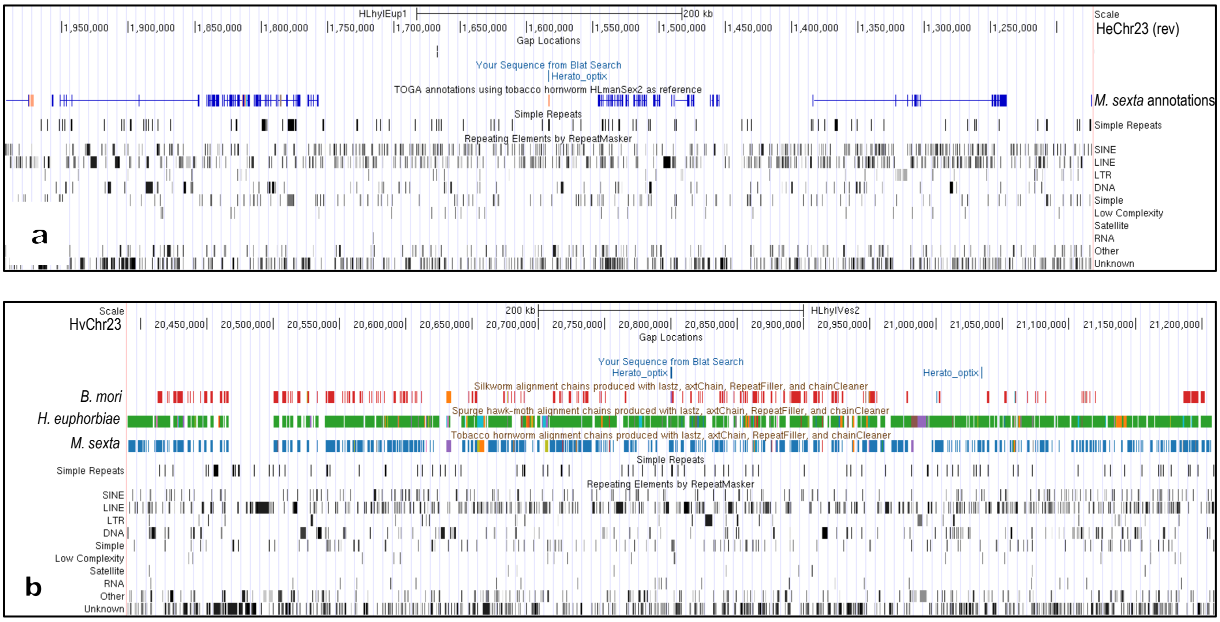


**BLAT result of *Heliconius optix* protein sequence (KC469894) from the genome browser on chromosome 23 in a) *Hyles euphorbiae* (reverse) and b) *Hyles vespertilio* in a 700 KB view.**

Zoomed out, the 700 KB view of the *Heliconius erato optix* protein sequence BLAT search reveals a high number of repeats in the vicinity of the stretch of gene exons in both *Hyles* species (*B. mori* and *M. sexta* are included in b) for comparison). The position, length, and type differ between species.

The annotation track (orange and bright blue) shows positions of exons annotated with TOGA, based on *M. sexta*.

Other sequence colors in b) are red (main scaffold of *B. mori*), green (main scaffold of *H. euphorbiae*), blue (main scaffold of *M. sexta*). Additional scaffolds are given yet another color to be able to distinguish which chains align which scaffold. The darker the shades of grey of the repeats, the closer they are to the repeat consensus sequence.

**Fig. S4**

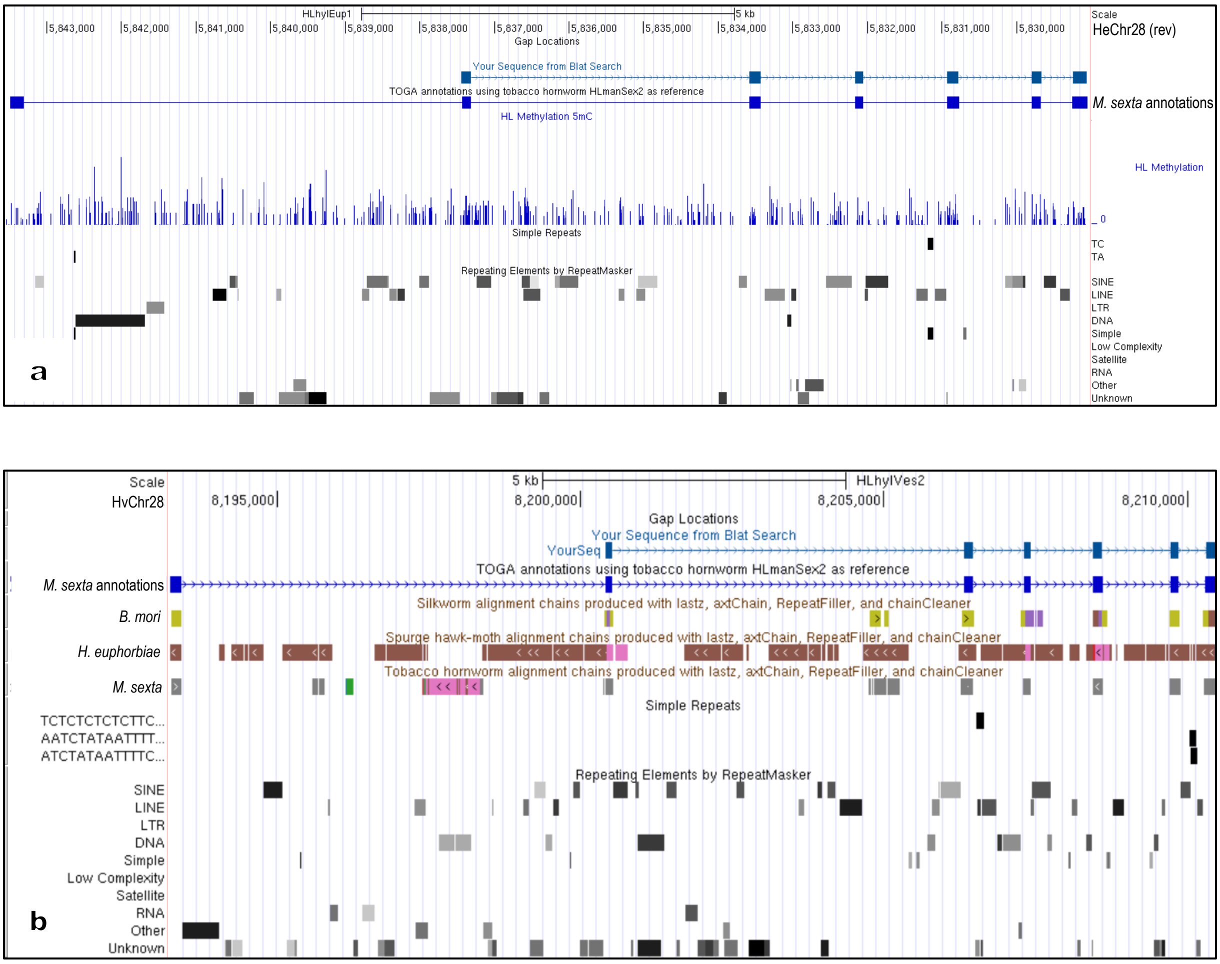


**BLAT result of the *Heliconius WntA* protein sequence (AFC75686) from the genome browser showing the *Hyles* genome alignment, annotation, and repeat occurrence in the vicinity in a) *H. euphorbiae* (reverse) and b) *H. vespertilio*.**

The *Heliconius himera WntA* amino acid sequence BLAT search resulted in a match corresponding to six exons on chromosome 28 in both *Hyles* species (*B. mori* and *M. sexta* are included in b) for comparison).

The annotation track (bright blue) shows positions of exons annotated with TOGA, based on *M. sexta*. The track “HL Methylation” in a) shows the *H. euphorbiae* methylation pattern analyzed, base modifications with probability < 50% are colored blue.

Other sequence colors in b) are light-green (main scaffold of *B. mori*), brown (main scaffold of *H. euphorbiae,* inverted) and grey (main scaffold of *M. sexta*). Additional scaffolds are given yet another color to be able to distinguish which chains align which scaffold. The darker the shades of grey of the repeats, the closer they are to the repeat consensus sequence.

**Fig. S5**

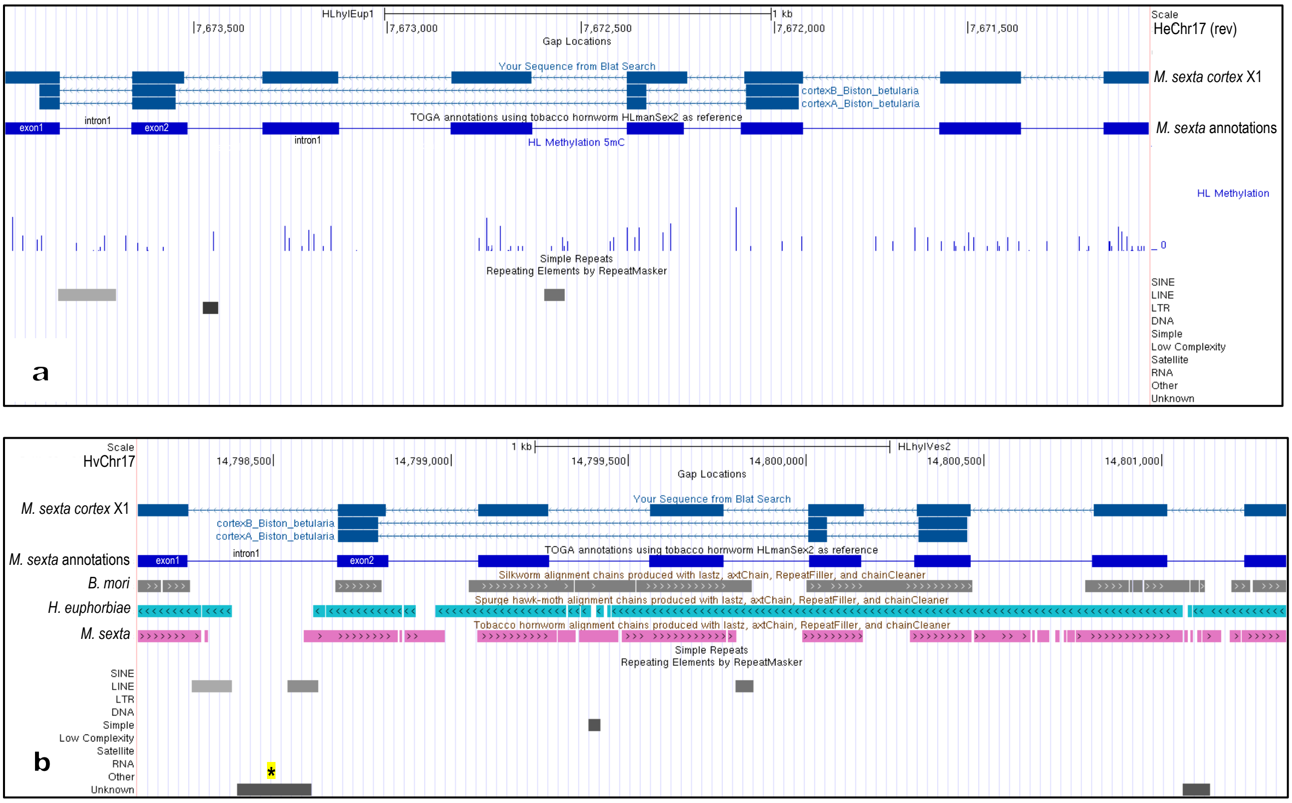


**BLAT result from the genome browser showing the exon/intron structures of the *Manduca sexta* and *Biston betularia cortex* protein (Table S2) on chromosome 17 in a) *Hyles euphorbiae* and b) *H. vespertilio*.**

The *M. sexta* cortex protein sequence BLAT search resulted in a match corresponding to eight exons on chromosome 17 in both *Hyles* species (*B. mori* and *M. sexta* are included in b) for comparison).

The annotation track (bright blue) shows positions of exons annotated with TOGA, based on *M. sexta*; exons 1, 2 and intron 1 are labelled for better orientation. The track "HL Methylation" in a) shows the *H. euphorbiae* methylation pattern analyzed, base modifications with probability < 50% are colored blue.

Other sequence colors in b) are grey (main scaffold of *B. mori*), blue-green (main scaffold of *H. euphorbiae*) and pink (main scaffold of *M. sexta*). The darker the shades of grey of the repeats, the closer they are to the repeat consensus sequence. The star (* highlighted in yellow) points to a stretch of (unknown) repeats within intron 1 unique for *H. vespertilio*.

**Fig. S6**

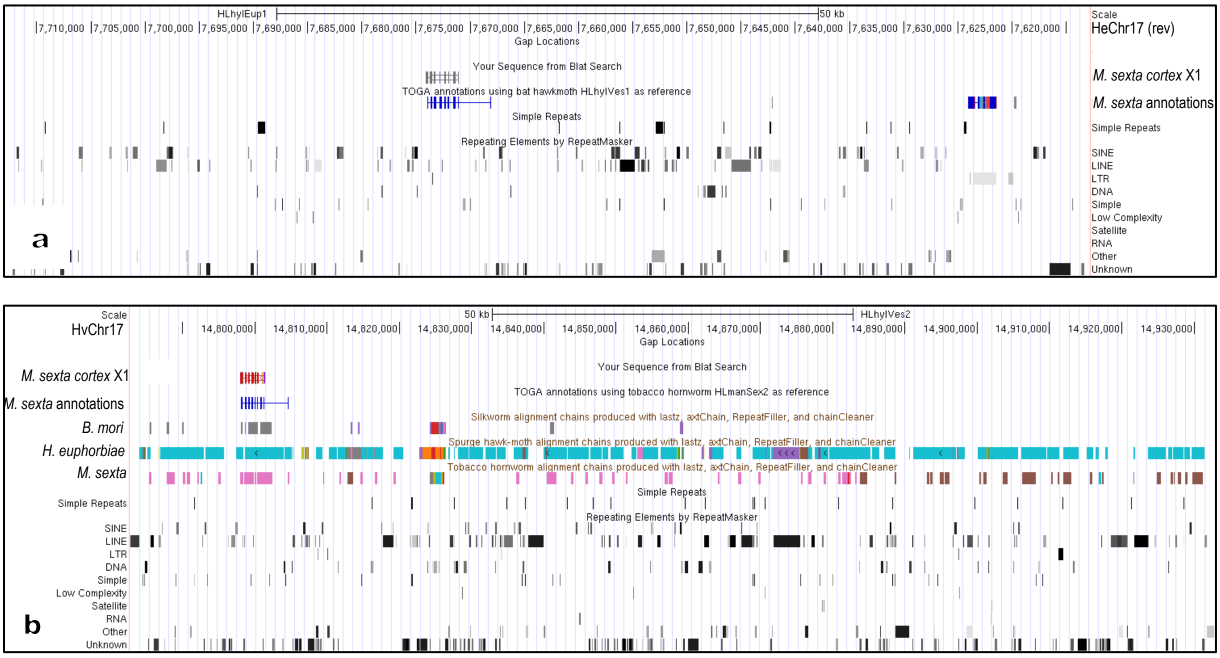


**BLAT result of *Manduca sexta* mRNA of *cortex* (Table S2) on chromosome 17 in a) *Hyles euphorbiae* and b) *Hyles vespertilio* in a 100 KB view.**

Zoomed out, the 100 KB view of the *M. sexta cortex* protein sequence BLAT search reveals a high number of repeats in the vicinity of the stretch of gene exons in both *Hyles* species (*B. mori* and *M. sexta* are included in b) for comparison). The position, length, and type differ between species.

Colors of the BLAT result in the genome browser are red if genome and query sequence have different bases, orange if query sequence has an insertion at this position, purple if query sequence extends beyond the end of the alignment, green if query sequence appears to have a polyA tail that is not aligned to the genome.

Other sequence colors in b) are grey (main scaffold of *B. mori*), blue-green (main scaffold of *H. euphorbiae*, inverted) and pink (main scaffold of *M. sexta*). Additional scaffolds are given a different color to be able to distinguish which chains align which scaffold. The darker the shades of grey of the repeats, the closer they are to the repeat consensus sequence.

**Fig. S7**

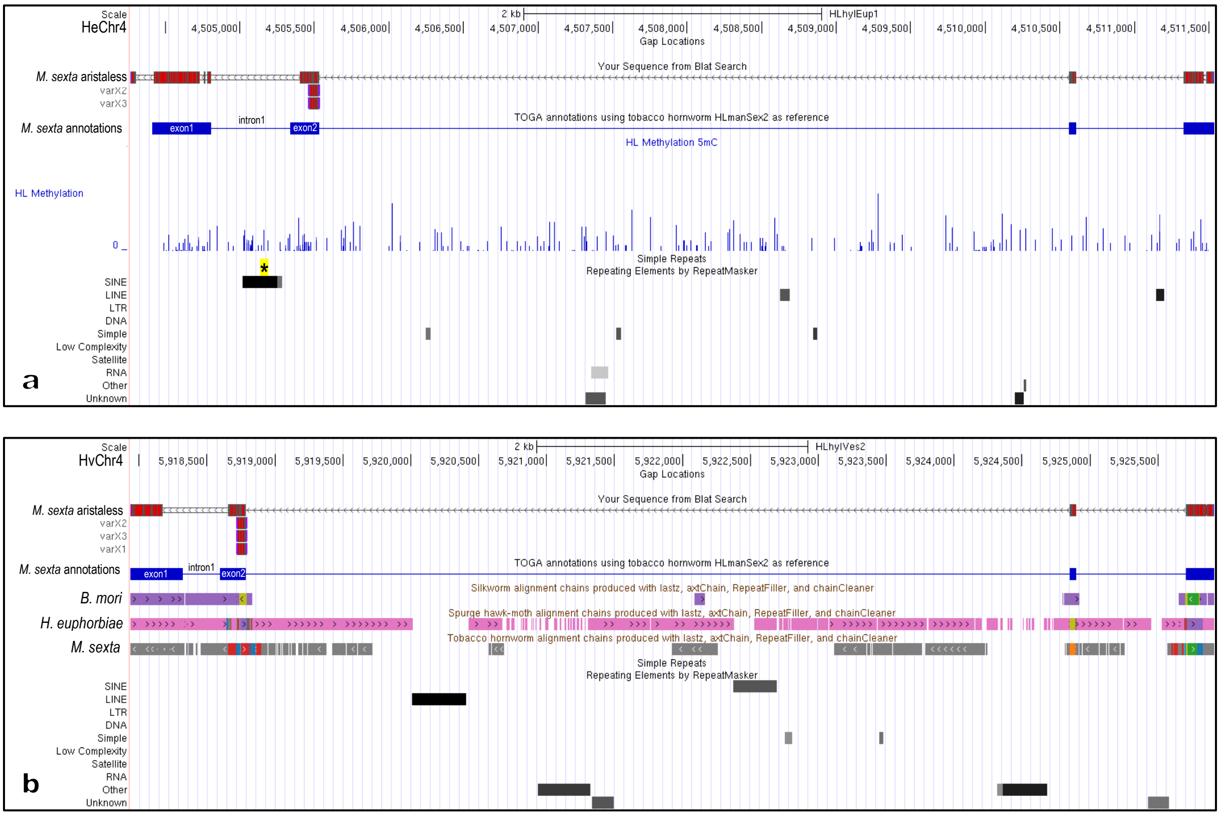


**BLAT result of *Manduca sexta* mRNAs of *aristaless* (Table 2) on chromosome 4 in a) *H. euphorbiae* and b) *H. vespertilio*.**

The *M. sexta aristaless* mRNA sequence BLAT search resulted in a match of several exons on chromosome 4 in both *Hyles* species (the stretch of the first 4 exons is shown; *B. mori* and *M. sexta* are included in b) for comparison).

Colors of the BLAT result in the genome browser are red if genome and query sequence have different bases, orange if query sequence has an insertion at this position, purple if query sequence extends beyond the end of the alignment, green if query sequence appears to have a polyA tail that is not aligned to the genome.

TOGA annotation using *M. sexta* is illustrated in bright blue, exons 1, 2 and intron 1 are labelled for better orientation in b).

Other sequence colors in b) are purple (main scaffold of *B. mori*), pink (main scaffold of *H. euphorbiae*) and grey (main scaffold of *M. sexta*). Additional scaffolds are given a different color to be able to distinguish which chains align which scaffold. The star (* highlighted in yellow) in a) points to a stretch of repeats within intron 1 that is unique for *H. euphorbiae*. The darker the shades of grey of the repeats, the closer they are to the repeat consensus sequence.

**Fig. S8**

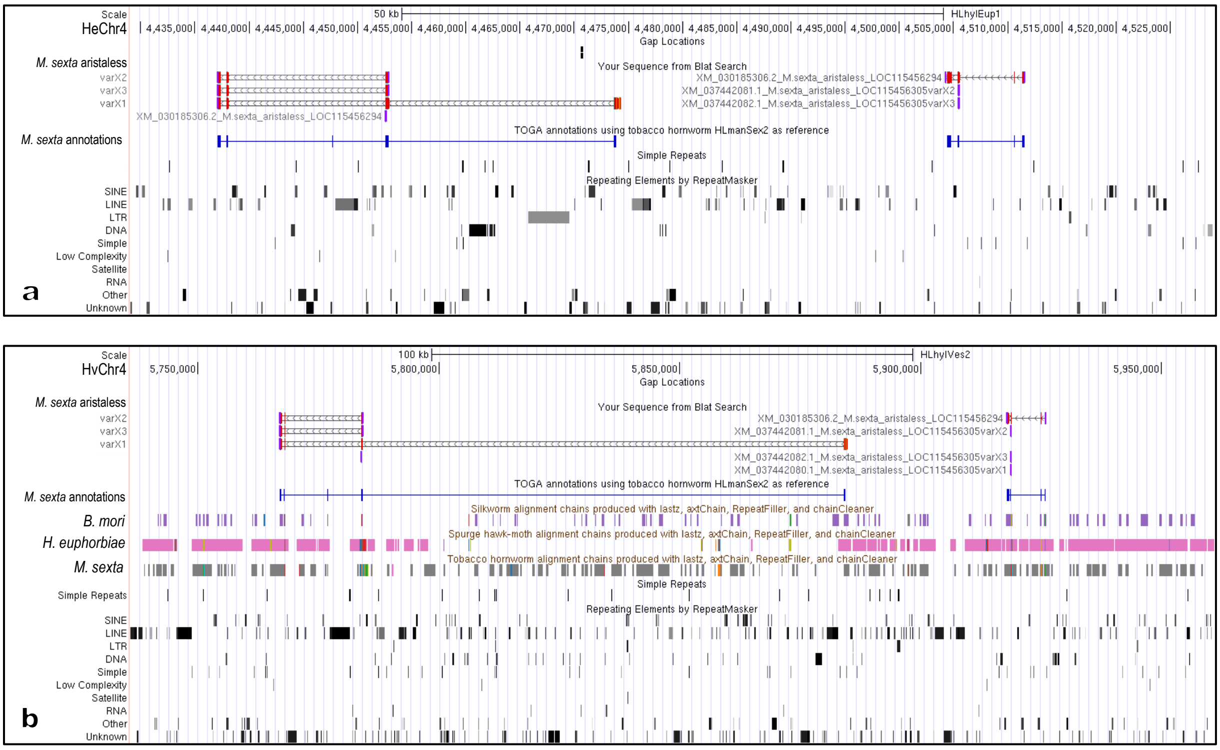


**BLAT result of *Manduca sexta* mRNAs of *aristaless* (Table 2) on chromosome 4 in a) *H. euphorbiae* 100K view, b) *H. vespertilio* 150 KB view (note the different scales).**

The *M. sexta aristaless* mRNA sequence BLAT search resulted in a match on chromosome 4 in both *Hyles* species, separated by 10-15 KB (*B. mori* and *M. sexta* are included in b) for comparison). Colors of the BLAT result in the genome browser are red if genome and query sequence have different bases, orange if query sequence has an insertion at this position, purple if query sequence extends beyond the end of the alignment, green if query sequence appears to have a polyA tail that is not aligned to the genome.

TOGA annotation using *M. sexta* is illustrated in bright blue.

Other sequence colors in b) are purple (main scaffold of *B. mori*), pink (main scaffold of *H. euphorbiae*) and grey (main scaffold of *M. sexta*). Additional scaffolds are given a different color to be able to distinguish which chains align which scaffold. The darker the shades of grey of the repeats, the closer they are to the repeat consensus sequence.

**Fig. S9**

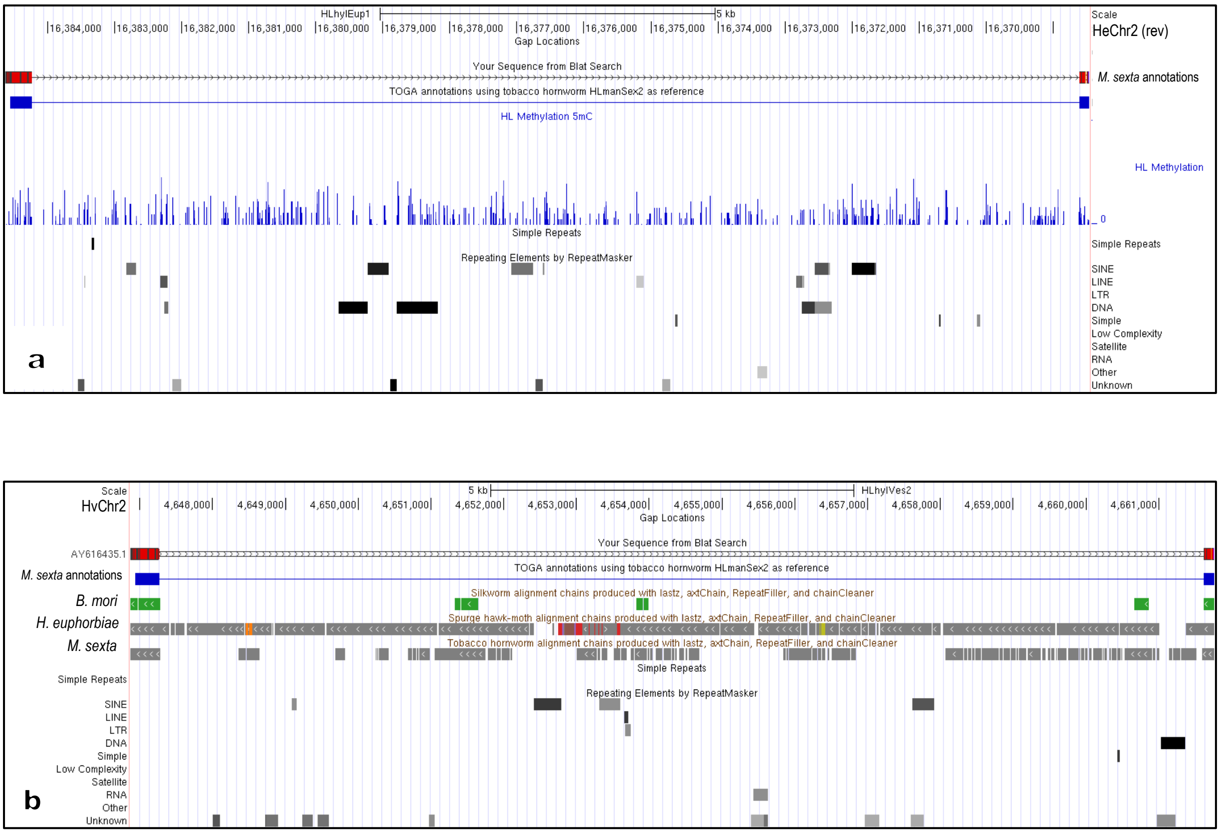


**BLAT result of *Manduca sexta* mRNA of *distal-less* (AY616435.1) on chromosome 2 in a) *Hyles euphorbiae* (inverted) and b) *Hyles vespertilio*.**

The *M. sexta distal-less* mRNA sequence BLAT search resulted in a match corresponding to two exons (TOGA annotation using *M. sexta* is shown in bright blue) on chromosome 2 in both *Hyles* species, separated by around 13 KB (*B. mori* and *M. sexta* are included in b) for comparison). Colors of the BLAT result in the genome browser are red if genome and query sequence have different bases, orange if query sequence has an insertion at this position, purple if query sequence extends beyond the end of the alignment, green if query sequence appears to have a polyA tail that is not aligned to the genome.

Other sequence colors in b) are green (main scaffold of *M. sexta*), grey (main scaffolds of *H. euphorbiae* & *B. mori* respectively). Additional scaffolds are given a different color to be able to distinguish which chains align which scaffold. The darker the shades of grey of the repeats, the closer they are to the repeat consensus sequence.

**Fig. S10**

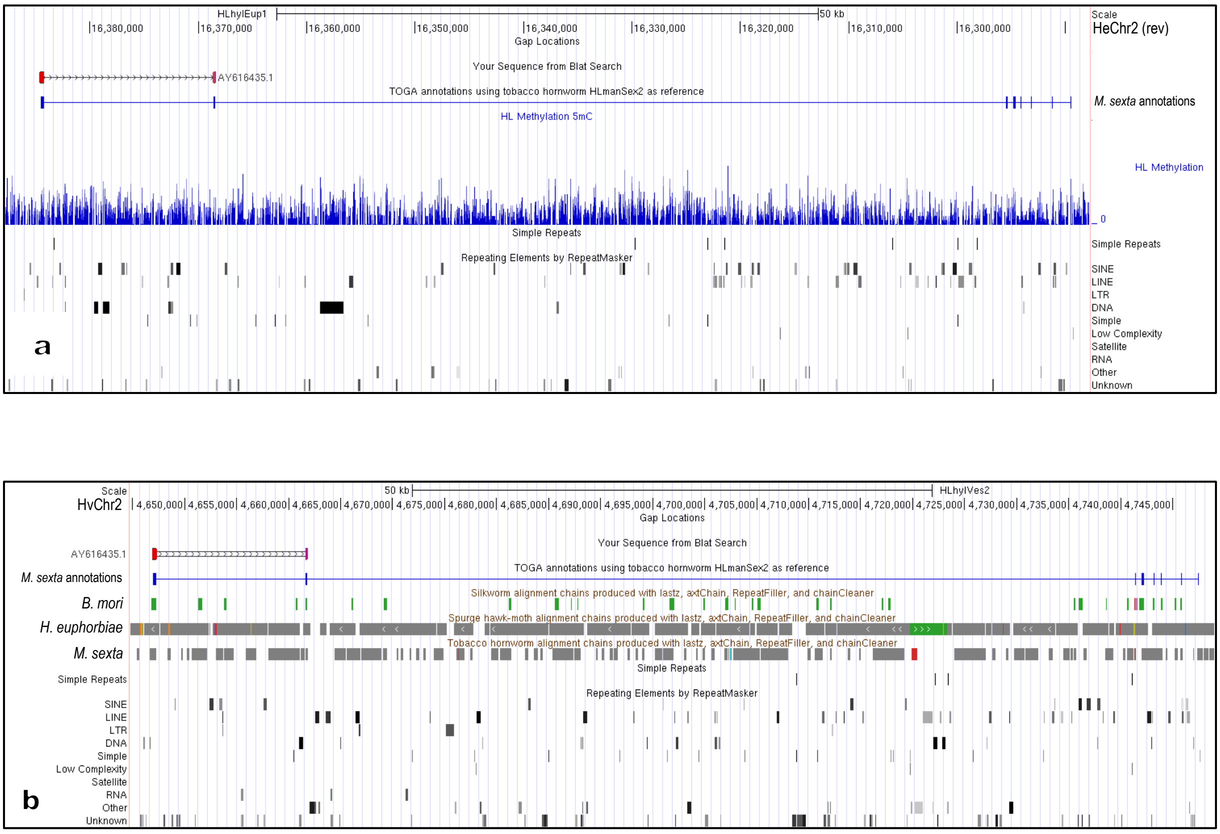


**BLAT result of *Manduca sexta* mRNA of *distal-less* (AY616435.1) on chromosome 2 in a) *Hyles euphorbiae* (inverted) and b) *Hyles vespertilio* in a 100 KB view.**

The *M. sexta distal-less* mRNA sequence BLAT search resulted in a match corresponding to two exons (TOGA annotation using *M. sexta* is shown in bright blue) on chromosome 2 in both *Hyles* species, separated by around 13 KB (*B. mori* and *M. sexta* are included in b) for comparison). Colors of the BLAT result in the genome browser are red if genome and query sequence have different bases, orange if query sequence has an insertion at this position, purple if query sequence extends beyond the end of the alignment, green if query sequence appears to have a polyA tail that is not aligned to the genome.

Other sequence colors in b) are green (main scaffold of *B. mori*), grey (main scaffolds of *H. euphorbiae* & *M. sexta* respectively). Additional scaffolds are given a different color to be able to distinguish which chains align which scaffold. The darker the shades of grey of the repeats, the closer they are to the repeat consensus sequence.
